# Subcellular reorganization upon phage infection reveals stepwise assembly of viral particles from membrane-associated precursors

**DOI:** 10.1101/2025.06.15.659807

**Authors:** Simon Corroyer-Dulmont, Audrey Labarde, Vojtěch Pražák, Lia Godinho, Chloé Masson, Pierre Legrand, Kay Grünewald, Paulo Tavares, Emmanuelle R.J. Quemin

## Abstract

Viruses are obligate intracellular parasites and viral infections lead to massive host cell rearrangement to support the rapid generation of progeny. Host take-over and remodelling include formation of viral-induced compartments for viral genome replication and/or assembly. While viruses infecting bacteria, bacteriophages or phages, have been extensively characterized *in vitro*, the molecular mechanisms underlying the viral cycle inside the crowded cytoplasm remain unclear. Here, we investigate the spatial reorganization of SPP1-infected bacteria under near-native conditions by electron cryo tomography. The most prominent feature is the formation of a large viral DNA (vDNA) compartment from which ribosomes are excluded. In SPP1 infection, there is no membrane nor proteinaceous shell surrounding these compartments. Also, we identified novel key intermediates in virus assembly: open precursors of procapsid lattice were found at the cytoplasmic membrane in a process that requires expression of the portal protein. Next, DNA-free procapsids relocate inside the vDNA compartment where vDNA is packed in a stepwise manner. Finally, DNA-filled capsids segregate to the periphery of the compartment for assembly completion and storage. Collectively, we provide comprehensive mechanistic insights into the complete viral assembly pathway of SPP1 directly *in cellula* and show how specific steps are coordinated inside the reorganized bacterial cell.

## Main text

Viruses are obligatory parasites that can infect bacteria, archaea or eukaryotes, and show co-evolution with their hosts. Several viral lineages are deeply rooted with common ancestry such as tailed bacterial viruses (bacteriophages or phages) together with herpesviruses^1^. During lytic infection, viruses massively hijack the resources of the host towards a rapid generation of progeny. This take-over promotes reorganization of the infected cell, frequently leading to formation of compartments dedicated to viral genome replication, progeny virion assembly and/or evasion from host defence systems^2^. Several strategies for viral-induced compartmentation are known: membrane-bound, confined by a protein cage, formed by liquid-liquid phase separation or result of polymer compaction (*e.g.*, DNA molecules) upon macromolecular crowding^3^.

In the case of bacteriophages, the main steps of viral multiplication were studied extensively *in vitro*. However, it remains largely unclear how well this represents the molecular processes actually taking place in the complex cytoplasmic environment inside infected cells. Recently, jumbo phages were shown to form a proteinaceous shell making a nucleus-like compartment, which physically separates replication and transcription of the viral genome from translation, viral particle assembly, and metabolic reactions inside the host bacterium^4^. This feature of giant jumbo phages (genome sizes above 200 kbp) distinguishes them from the vast majority of tailed viruses that have smaller genomes, most frequently in the 30-60 kbp size range^5^. Notably, infection by siphoviruses lambda^6^ and SPP1^7^ also lead to the formation of a viral DNA (vDNA) compartment that is localized asymmetrically within the bacterial host cell. This compartment appears to play a pivotal role in phage DNA transactions (*i.e*., replication, recombination and encapsidation) but does not protect the viral genome against bacterial defence mechanisms^8^. Furthermore, in the case of SPP1, DNA-filled viral particles are stored in so-called warehouses found at the periphery of the vDNA compartment^7^.

Here, we investigate the spatial reorganization of the host bacterial cell induced upon infection by the bacteriophage SPP1 and characterize key individual steps of the viral particle assembly pathway *in cellula* under near-native conditions. Using cellular electron cryo tomography (cryoET), we could identify assembly intermediates previously undetected in *in vitro* studies: formation of procapsid precursors at the cellular membrane as well as subsequent packaging of vDNA into the procapsid structure. Importantly, we also find that these viral molecular processes correlate with a specific spatial partitioning within the rearranged infected cell either at the plasma membrane, inside the vDNA compartment or confined to its periphery. Altogether, we provide comprehensive mechanistic insights into the complete assembly pathway from membrane-associated precursors of procapsids to DNA-filled head-and-tail capsids directly in the bacterial host cell.

### Electron cryo tomography applied to study SPP1-infected bacteria

We developed an optimized sample preparation workflow to follow SPP1 infection *in cellula* (Extended Data Fig. 1) and measure its impact on the remodelled infected bacteria under near-native conditions^9^. The well-characterized siphovirus SPP1 infects the Gram-positive soil bacterium *Bacillus subtilis*^10^. Under optimal infection conditions, SPP1 DNA is found inside bacterial cells within 3 min post infection (p.i.)^11^. Transcription and replication of the vDNA genome are initiated rapidly afterwards^12^, while expression of late genes coding for morphogenetic proteins starts at 10 to 12 min p.i.^11^. From 30 min p.i. onwards, infectious viral particles are formed and then released by cell lysis^11^. In order to visualize all states of virus assembly, we generated specimens of SPP1-infected *B. subtilis* cells at 15 and 25 min p.i. by plunge-freezing (see Online Methods). Then, thin lamellae (*i.e.*, sections of ∼100 nm in thickness) were micromachined under cryogenic conditions using a scanning electron microscope equipped with a focused ion beam (cryoFIB-SEM) (Extended Data Fig. 1)^13^. Each lamella (∼12 µm in width) contained cross-sections of several bacteria (Extended Data Fig. 1f) infected either with wild type (*wt*) or mutant SPP1 phages that were investigated by cryoET (Extended Data Fig. 2).

### SPP1 infection leads to extensive remodelling of *B. subtilis* cells and formation of a prominent vDNA compartment

*B. subtilis* cells were imaged before and after infection with SPP1 using cryoET (Fig. 1; Extended Data Fig. 3). Tomograms of uninfected control cells display a homogeneous distribution of ribosomes in the cytoplasm (Fig. 1a; Extended Data Fig. 3a; Supplementary Movie 1), which starkly contrasts with infected cells where ribosomes are mostly excluded from a large area of the cytoplasm (Fig. 1b; Extended Data Fig. 3b; Supplementary Movie 2). There, a prominent compartment is formed that contains the vDNA (Extended Data Fig. 3b-h). Indeed, fluorescence microscopy of ribosomal proteins L1 and S2 showed that reorganization of the cytoplasmic space correlated with the exclusion of ribosomes from the phage vDNA compartment, and late in infection with the formation of adjacent warehouses where mature infectious virions are stored (Extended Data Fig. 4). Notably, these two types of viral-induced compartments are neither delineated by a lipid bilayer nor by a proteinaceous shell (Fig. 1; Extended Data Fig. 3).

**Fig. 1.**
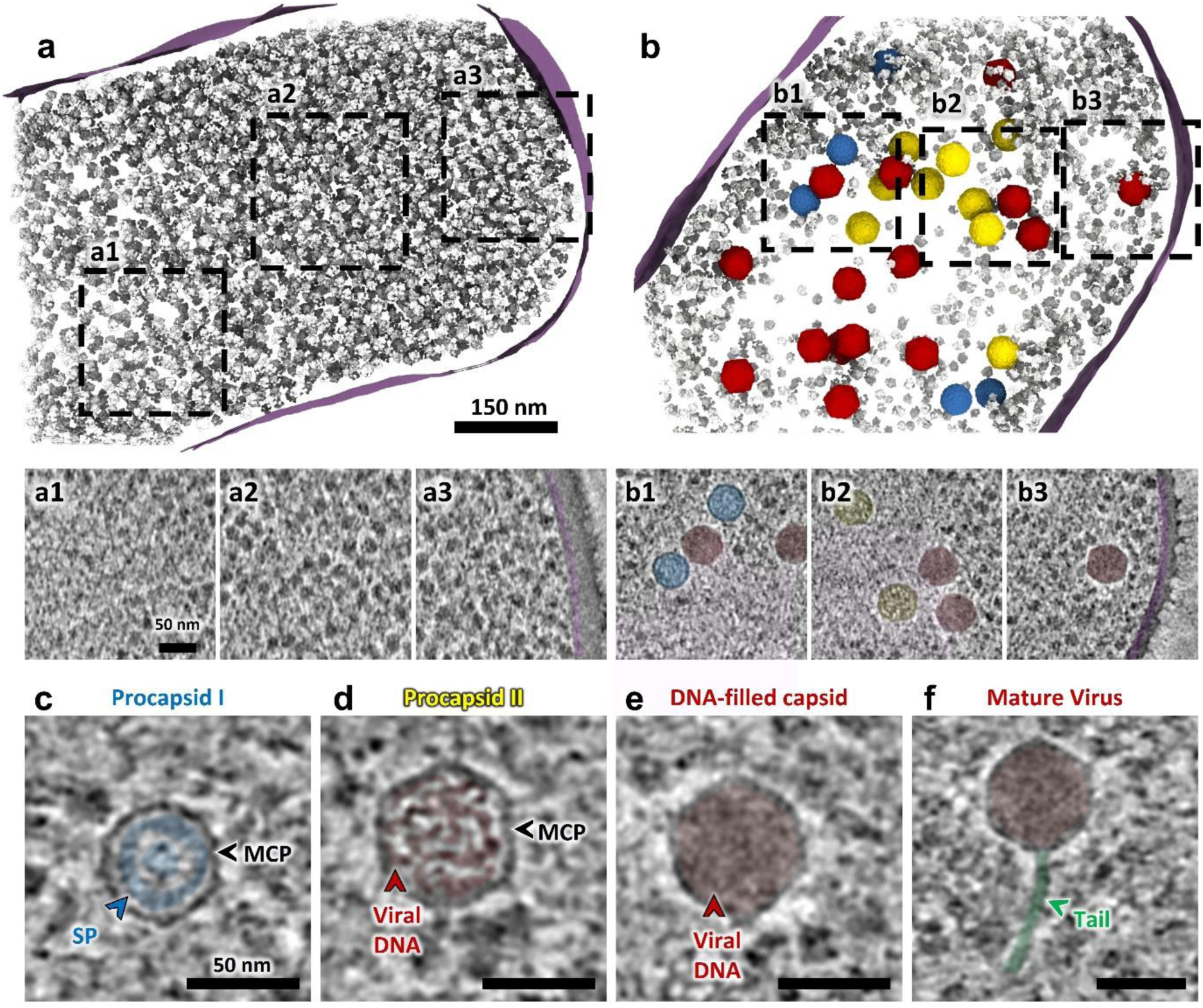
Analysis of SPP1-induced intra-bacterial reorganization and compartmentalization of viral processes observed *in cellula*. **a**. Segmentation rendering and back-plotting of a representative tomogram (Supplementary Movie 1) of non-infected *B. subtilis* exhibiting the cellular organization with the bacterial cytoplasm filled with ribosomes (grey) and surrounded by the cell membrane (magenta). Zoomed-in insets of tomogram regions indicated by dashed lines in the segmentation rendering show details of a small region with bacterial DNA (**a1**) while the rest of the cytoplasm appears homogeneously filled with ribosomes (**a2**, **a3**). **b**. Segmentation rendering and back-plotting of a representative tomogram (Supplementary Movie 2) acquired on *B. subtilis* infected by SPP1 wild type at 25 min p.i. in which a region of exclusion of ribosomes is clearly visible. Capsids at different steps of the DNA packaging process – procapsid I (blue), and procapsid II (yellow), are found in this region (detail in **b1, b2**). DNA-filled capsid (red) are present inside the vDNA compartment and at the periphery while mature virions localize primarly at the vDNA compartment periphery (**b3**) (see distribution in Extended Data Fig. 2). The region of ribosome exclusion is filled with filaments that most likely represent the viral DNA (pink in **b1** and **b2**). **c-f**. Insets of slices from representative tomograms displaying the assembly intermediates of SPP1: procapsid I (**c**), procapsid II partially filled with DNA (**d**), DNA-filled capsid without tail (**e**), and a mature virus with a tail (**f**). Colouring of features of interest: (**c**) scaffolding protein (SP) in blue, major capsid protein (MCP) is indicated by black arrowheads; (**d-f**) viral DNA in red; (**f**) tail in green.

### Spatial coordination of the different stages of viral particle assembly inside the host bacterial cell

Direct visualization of individual phage particles *in cellula* using cryoET overcomes a long-standing bottleneck in the field posed by the inability to image those structures by conventional electron microscopy (EM) approaches after sample preparation on chemically fixed bacteria following infection^7^. In *wt* infections, we observed that the majority of viral particles can be assigned to several intermediates of the capsid assembly process, namely procapsid I, procapsid II, DNA-filled capsid or mature virus (Fig. 1c-f). Structures of three of those assembly states have previously been determined from purified particles by cryoEM single particle analysis^14^. Accordingly, procapsids I are spherically-shaped icosahedra, 55 nm in diameter, with the scaffolding proteins (SP, highlighted in blue in Fig. 1c) in the lumen. Procapsids II exhibit an angular icosahedral structure with a diameter of 65 nm and were here frequently filled with various amounts of DNA (Fig. 1d). Finally, DNA-filled capsids with similar dimensions to procapsids II (Fig. 1e) can be found at the periphery of the vDNA compartment, together with mature virions attached to a long flexible tail (Fig. 1f; Extended Data Fig. 2a).

### Precursors of SPP1 procapsids assemble at the bacterial cell membrane

Notably, partial SPP1 procapsid I-like structures were consistently associated to the inner side of the bacterial cell membrane (Fig. 2; Extended Data Fig. 5). We consider these incomplete procapsids as early precursors of SPP1 assembly. They account for more than 10% of viral particles observed in our tomograms at 15 min p.i. and about 3% at 25 min p.i. (Extended Data Fig. 2c). Based on the statistical distribution of precursors with different degrees of completeness, it appears that initial capsomere assembly is rapid but subsequent procapsid completion and membrane dissociation might be the limiting steps (Fig. 2i). The stages of precursor assembly observed vary from about one third of an opened sphere (Fig. 2a,e; Extended Data Fig. 5a), up to closed procapsid I-like shells still bound to the membrane (Fig. 2d,h; Extended Data Fig. 5d). The diameter of those precursors (52 nm on average, SD=0.7 nm, n=20) as well as the curvature of the thick outer capsid lattice are similar to the unexpanded lattice of procapsids I observed *in cellula* (this work) and in the high-resolution structure previously determined *in vitro*^14^ (Extended Data Fig. 6). The lumen of precursors consists of one or two layers of, presumably, internal scaffolding proteins that serve for capsomere assembly (SP, highlighted in blue in Fig. 2e-h). A pattern of radial striations is indeed observed within the outer shell, giving it the appearance of individual concentric rods (Fig. 2a-h,j). Subvolume averaging was performed on procapsids I found in our tomograms in order to determine the symmetry of the scaffolding layers with alignment focusing either on the icosahedral shell or on the protein scaffold (see Online Methods). No structural detail was discernible for the scaffolding proteins independent of whether C12 symmetry was applied or not, implying that the scaffold does not follow a defined regular icosahedral organization in these procapsids but a different and possibly asymmetrical shell packing (Extended Data Fig. 6a,b).

**Fig. 2.**
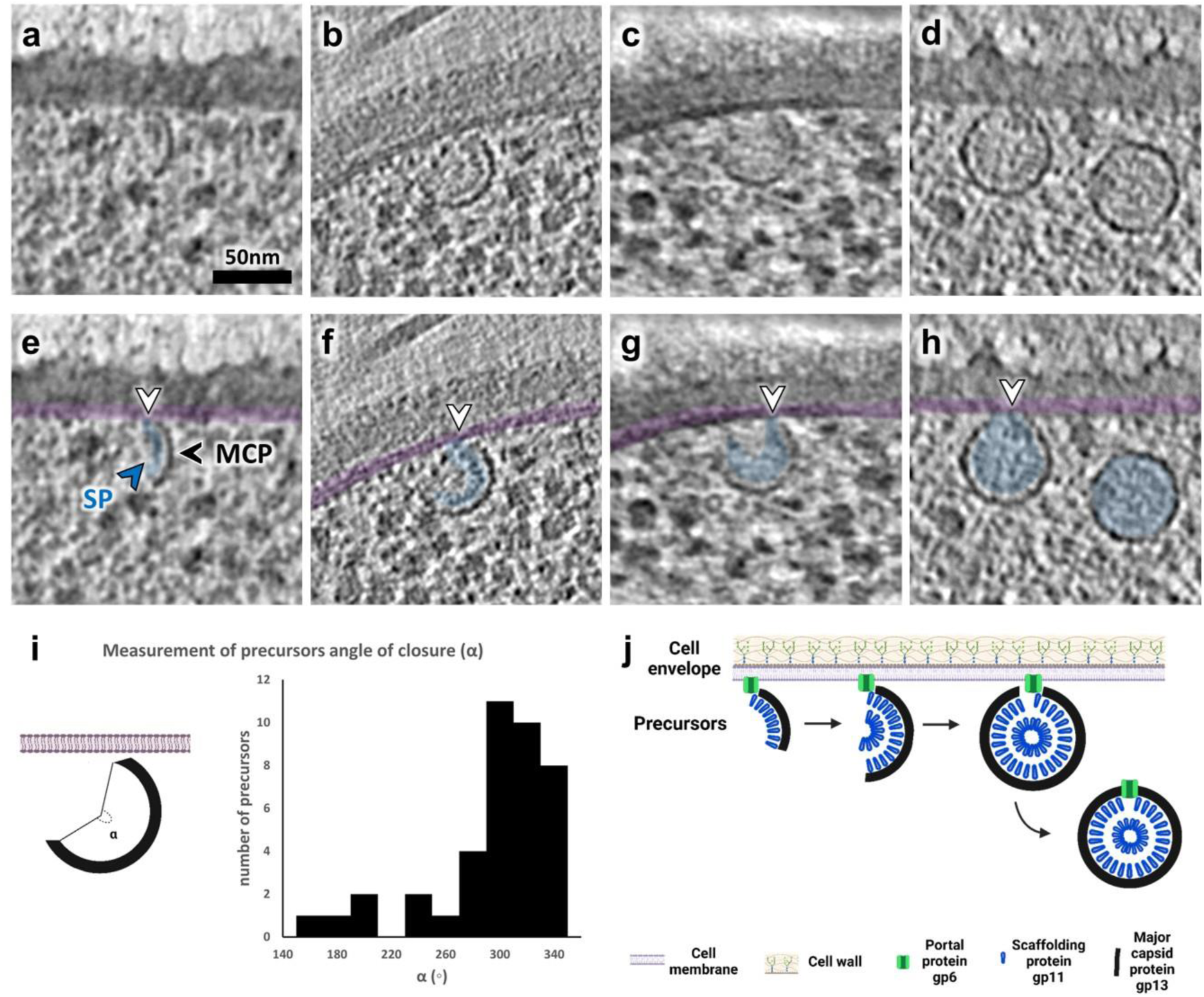
Stepwise assembly of SPP1 procapsids is initiated by formation of precursors at the inner side of the bacterial cell membrane. **a-d**. Zoomed-in insets of slices from tomograms of *B. subtilis* infected by SPP1 displaying procapsid precursors associated to the inner side of the bacterial cell membrane at different stages of procapsid lattice growth. Notably, the complete procapsid I particle is found detached from the cell membrane in (**d)**. Serial slices through the tomograms are provided in Extended Data Fig. 5. **e-h**. Transparent overlay colouring of slices shown in (**a-d)** to highlight the internal scaffolding protein (SP; in blue; blue arrowhead) visible in precursors and in procapsids I. The major capsid protein (MCP; black arrowhead) is also annotated, and the cell membrane is overlaid in magenta. White arrowheads indicate connections between the precursors and the cell membrane. **i**. Distribution of assembly intermediates of procapsid precursors found in tomograms according to their angle of closure (α) defined as in the cartoon on the left. **j**. Model of assembly of procapsid precursors initiated at the membrane-attached portal protein gp6 (green) followed by curvilinear co-assembly of scaffolding (gp11, blue) and major capsid (gp13, black) proteins.

### Role of the portal protein in membrane-associated procapsid assembly

All capsid precursors observed were in close proximity to the bacterial cell membrane and connections are sometimes visible between the inner side of the cell membrane and the capsid lattice being assembled (Fig. 2a-h; Extended Data Fig. 5). These suggest that the host membrane serves as an anchor point for nucleation of procapsid assembly initiation. We hence hypothesized that the portal protein gp6 could nucleate SPP1 procapsid assembly at the bacterial membrane. Consistent with this hypothesis, it was already reported that gp6 is soluble in solution^15^ but can also form nanopores in membranes under certain conditions *in vitro*^16^. In order to probe the localization of isolated SPP1 portal proteins *in cellula*, we engineered a *B. subtilis* strain producing gp6 whose carboxyl terminus was fused to the fluorescent protein mCitrine (see Online Methods). Gp6-mCitrine is functional for SPP1 assembly as assessed in a trans-complementation assay (Extended Data Fig. 7). These cells were infected by a gp6 deletion mutant strain (SPP1*gp6^-^*), ensuring that the sole source of portals for incorporation in capsids is the bacterial-encoded gp6-mCitrine. Epifluorescence microscopy observations showed a weak punctate fluorescence distribution in non-infected bacteria (Fig. 3a). We then used total internal reflection fluorescence (TIRF) microscopy to assess the dynamics of gp6-mCitrine at different stages of infection^17^. Early in infection, puncta of gp6-mCitrine are highly dynamic, but become immobilised in larger clusters at 23 min p.i., which coincides with viral particle assembly. Interestingly, these fluorescent foci localised to the periphery of the vDNA compartment (Fig. 3a,b; Supplementary Movies 3 and 4). We conclude that gp6-mCitrine proteins are mostly incorporated in viral procapsids that mature into virions and are subsequently stored in immobile warehouse compartments that were previously characterized^7^.

**Fig. 3.**
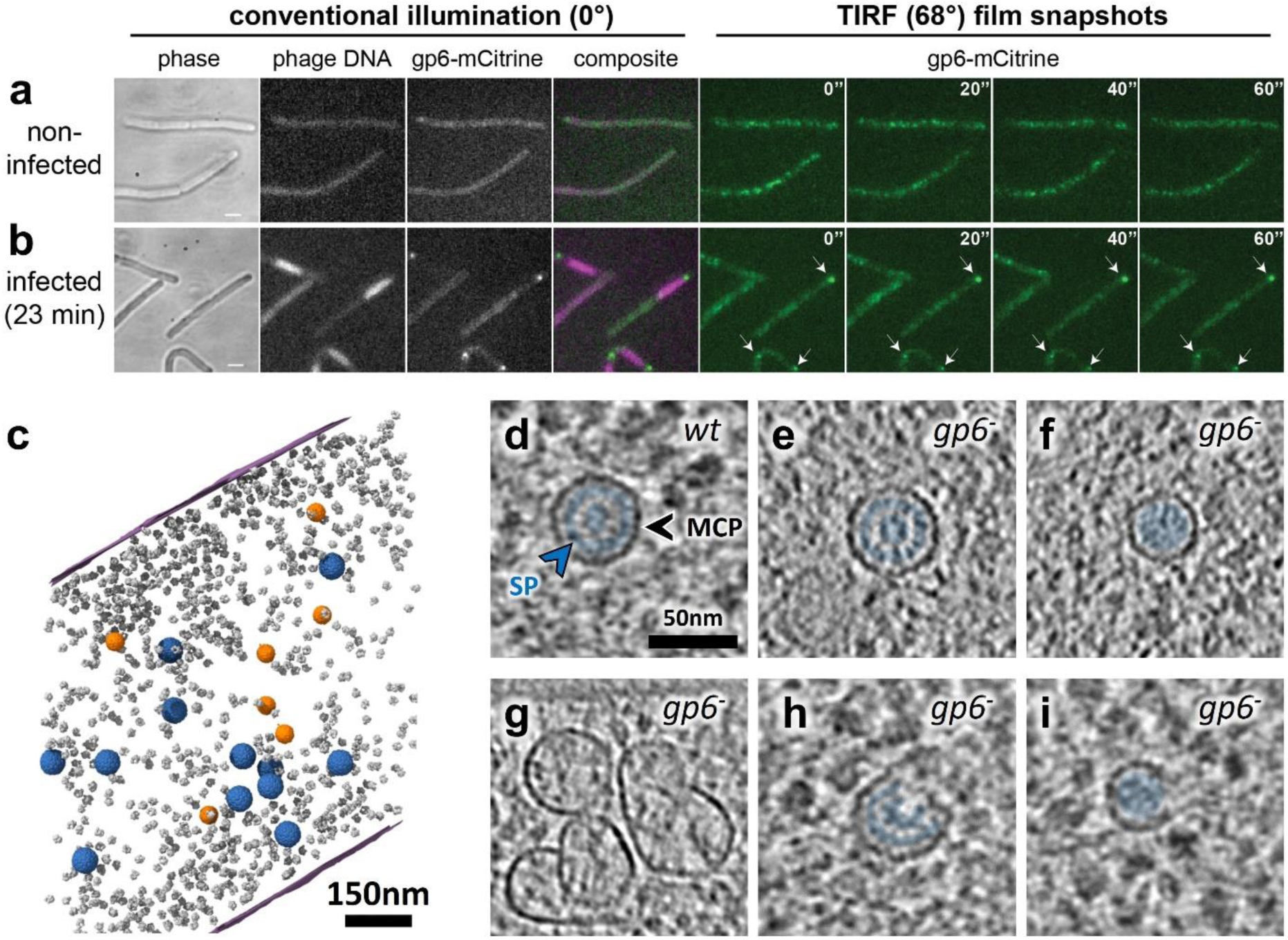
Localization of the SPP1 portal protein inside the bacterial cell and its role in membrane-associated assembly of procapsid precursors. **a**. Localization of the SPP1 portal protein gp6 fused to mCitrine (gp6-mCitrine) in non-infected bacterial cells imaged by phase contrast (phase) and fluorescence. Fluorescence imaging using conventional illumination (left panels; illumination at a 0° angle) and TIRF time-lapse (right panels; illumination at a 68° angle). TIRF film snapshots show the high mobility of gp6-mCitrine foci in the peri-membrane region (right) (Supplementary Movie 3; time-lapse acquisition every 2 s during 1 min). **b**. Localization of gp6-mCitrine and of phage DNA labelled by LacI-mCherry in cells infected at 23 min p.i. by SPP1*lacO64gp6^-^* that do not produce the portal protein gp6. Infected bacterial cells were imaged as in (**a**). White arrows show immobile foci of gp6-mCitrine in the TIRF time-lapse (Supplementary Movie 4). **c**. Segmentation rendering and back-plotting of a representative tomogram of *B. subtilis* infected by SPP1*gp6^-^* (Supplementary Movie 5). Procapsids I (blue) and smaller procapsid-like structures (∼40 nm in diameter; n = 111; orange) are found mostly in the vDNA compartment defined by ribosome-exclusion. Ribosomes are shown in grey and the bacterial cell membrane in magenta. **d-i**. Zoomed-in insets of tomogram slices depicting procapsids I in *B. subtilis* infected by SPP1 wild type (*wt*) (**d**) and SPP1*gp6^-^*(*gp6^-^*) (**e**). Different structures assembled in absence of the portal protein gp6 are also shown: small procapsid-like assembly intermediates not associated with the membrane (**f**), aberrant capsid-like material with various shapes and sizes that lack a clear inner scaffolding structure (**g**), and assembly intermediates of either procapsid-like structure (**h**) or of a small procapsid (**i**). The scaffolding protein (SP; blue arrowhead) is annotated in (**d**) and coloured (blue) when present (**d-f,h,i**). The major capsid protein (MCP; black arrowhead) forming the capsid shell is also indicated in (**d**).

We next investigated whether procapsid assembly occurs at the membrane in absence of the portal protein. In *B. subtilis* cells infected with SPP1*gp6^-^*, meaning conditions under which the portal is not produced (Fig. 3c; Extended Data Fig. 3c; Supplementary Movie 5), we found that only procapsid I-like particles (Fig. 3e), smaller procapsids (Fig. 3f) and some aberrant particles (Fig. 3g) were present. These observations are consistent with former studies showing that the portal protein is necessary for faithful size determination of SPP1 procapsids^18^. Strikingly, no procapsid assembly intermediates were observed to be associated with the cell membrane in the 58 tomograms analysed (Extended Data Fig. 2a). We hence hypothesize that the incomplete, open particles rarely spotted in the cytoplasm and in the vDNA compartment (n=5) represent intermediates of an alternative, gp6-independent assembly pathway under these specific conditions (*e.g*., Fig. 3h,i). This previously unsuspected requirement of gp6 for localization of procapsid precursors at the inner side of the bacterial cell membrane strongly suggests that it plays a key role in regulating and driving the early stages of SPP1 procapsid assembly.

Such finely tuned mechanism would favour in particular the incorporation of one portal at the initiation of assembly, prior to the icosahedral lattice polymerization step.

### Concomitant DNA packaging and capsid expansion occur after relocalization of procapsids to the vDNA compartment

To determine the spatial distribution of procapsids in absence of any vDNA packaging events, we imaged bacteria infected by SPP1 conditional mutants defective for expression of the viral terminase components TerS or TerL (gp1 and gp2 in SPP1, respectively; see Online Methods). TerS is known to selectively recognize the phage genome and to recruit TerL that docks the vDNA at the procapsid portal vertex, allowing packaging to proceed through the portal protein channel^19,20^. In SPP1*gp1^-^* and SPP1*gp2^-^* infections, capsid precursors are found associated to the membrane and procapsids I in the vDNA compartment, in a manner similar to *wt* virus infection (Fig. 4a; Extended Data Fig. 3d,e; Supplementary Movie 6). Hence, detachment of fully assembled procapsids I from the membrane and their relocation to the vDNA compartment occur independently of subsequent vDNA encapsidation events (Extended Data Fig. 2). Notably, procapsid-like structures without portal assembled during SPP1*gp6^-^*infection are also found in the vDNA compartment (blue in Fig. 3c; Extended Data Fig. 3c) demonstrating that their specific targeting to the compartment is independent of the portal, and likely relying on the procapsid icosahedral lattice properties.

**Fig. 4.**
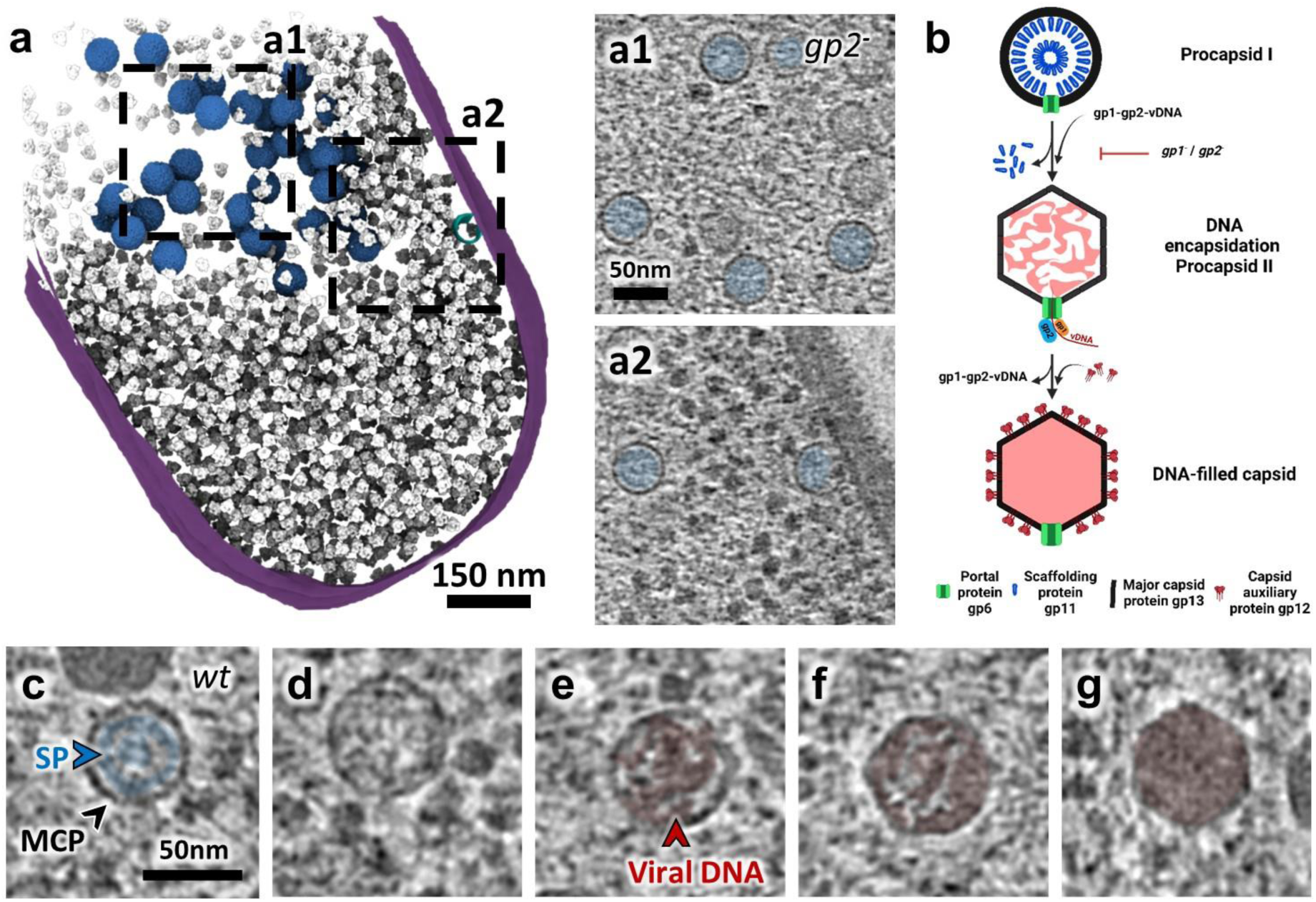
Viral DNA packaging and capsid expansion mediate the transition from procapsid I to procapsid II in the vDNA compartment. **a**. Segmentation rendering and back-plotting of a representative tomogram of *B. subtilis* infected by SPP1lacO64gp2-(*gp2^-^*) (Supplementary Movie 6), a mutant defective in viral DNA packaging. The bacterial cell membrane is shown in magenta, back-plotted ribosomes in grey, the procapsid precursor in cyan, and procapsids I in blue. Zoomed-in insets of regions indicated by dashed lines in the segmentation rendering show details of the vDNA compartment interior (**a1**) and periphery with a precursor at the cellular membrane **(a2)**. **b**. Schematic representation of viral DNA packaging steps during SPP1 assembly including procapsid I, procapsid II and DNA-filled capsid as well as concomitant steps of capsid expansion and release of the scaffolding proteins. **c-g**. Zoomed-in insets of slices from tomograms of *B. subtilis* infected by SPP1 wild type (*wt*) displaying intermediates of viral DNA genome packaging inside the vDNA compartment. **c.** Detail of procapsid I with the arrangement of the scaffolding protein (SP; blue arrowhead) gp11 (blue) visible on the inside as radial striations. The major capsid protein (MCP; black arrowhead), gp13, is also indicated. **d-f.** Details of individual procapsids II after capsid expansion containing increasing amount of viral DNA inside (red arrowhead and colouring) as packaging progresses (left to right). **g.** Detail of a DNA-filled capsid showing homogeneous density of the internal content, most likely representing the final stage of viral genome packaging.

The transition from procapsid I to procapsid II is marked by capsid expansion and release of the scaffolding proteins^14^. Procapsids II were present neither in SPP1*gp1^-^* nor in SPP1*gp2^-^*infections (Fig. 4a; Extended Data Fig. 2a; Extended Data Fig. 3d,e) showing that vDNA packaging is essential to trigger expansion of the procapsid lattice *in cellula* (Fig. 4c-g). Nevertheless, in a previous study, expanded procapsids II were found in extracts of lysed bacteria from a SPP1*gp1^-^* infection^14^. Those procapsids II did not contain scaffolding proteins associated to capsid hexamers but the ones bound to the vertex pentamers were still present. Given our new findings, it seems that the disruption of the cellular environment by lysis was the trigger for procapsid I to procapsid II transition in absence of vDNA packaging. This discrepancy to earlier findings highlights the limitations of relying uniquely on *in vitro* structural biology to investigate such complex molecular processes. In SPP1*wt* infection, procapsids II, compared to procapsids I, display a larger, more angular shape similar to the structures determined *in vitro*^14^. In snapshots of vDNA packaging found in tomograms, the genome appears as thin threads inside procapsids II, which all have the same dimensions (63 nm in diameter ±1.1 nm; n=37). Altogether, our findings suggest that expansion is essentially a single transition event triggered by vDNA packaging initiation. Different steps of genome encapsidation were visible from nearly empty to almost full icosahedral capsids (Fig. 4d-g). The viral genome appears to occupy the overall internal procapsid II space adopting progressively a more compact organization and no common or regular DNA encapsidation pattern was observed inside those packaging intermediates (Extended Data Fig. 8).

### Segregation of DNA-filled capsids to the periphery of the vDNA compartment and binding to pre-assembled tails

We noticed that procapsids I and II are confined at 97% (n=1478) and 94% (n=225), respectively, to the vDNA compartment consistent with the localization of vDNA encapsidation events mediating the transition between the two procapsid stages (Extended Data Fig 2a). In contrast, 40% of DNA-filled capsids (n=604) distinguishable by their electron dense inner content (Figs. 1e, 4g) and the vast majority of tailed virions, 84% (n=113), were located outside of this compartment, mostly at its periphery (Fig. 5; Extended Data Fig. 2a). We conclude that capsids depart from the vDNA compartment after packaging of the phage genome and that tail attachment to DNA-filled capsids takes place outside of the vDNA compartment but adjacent to it (Extended Data Fig. 9a,b).

**Fig. 5.**
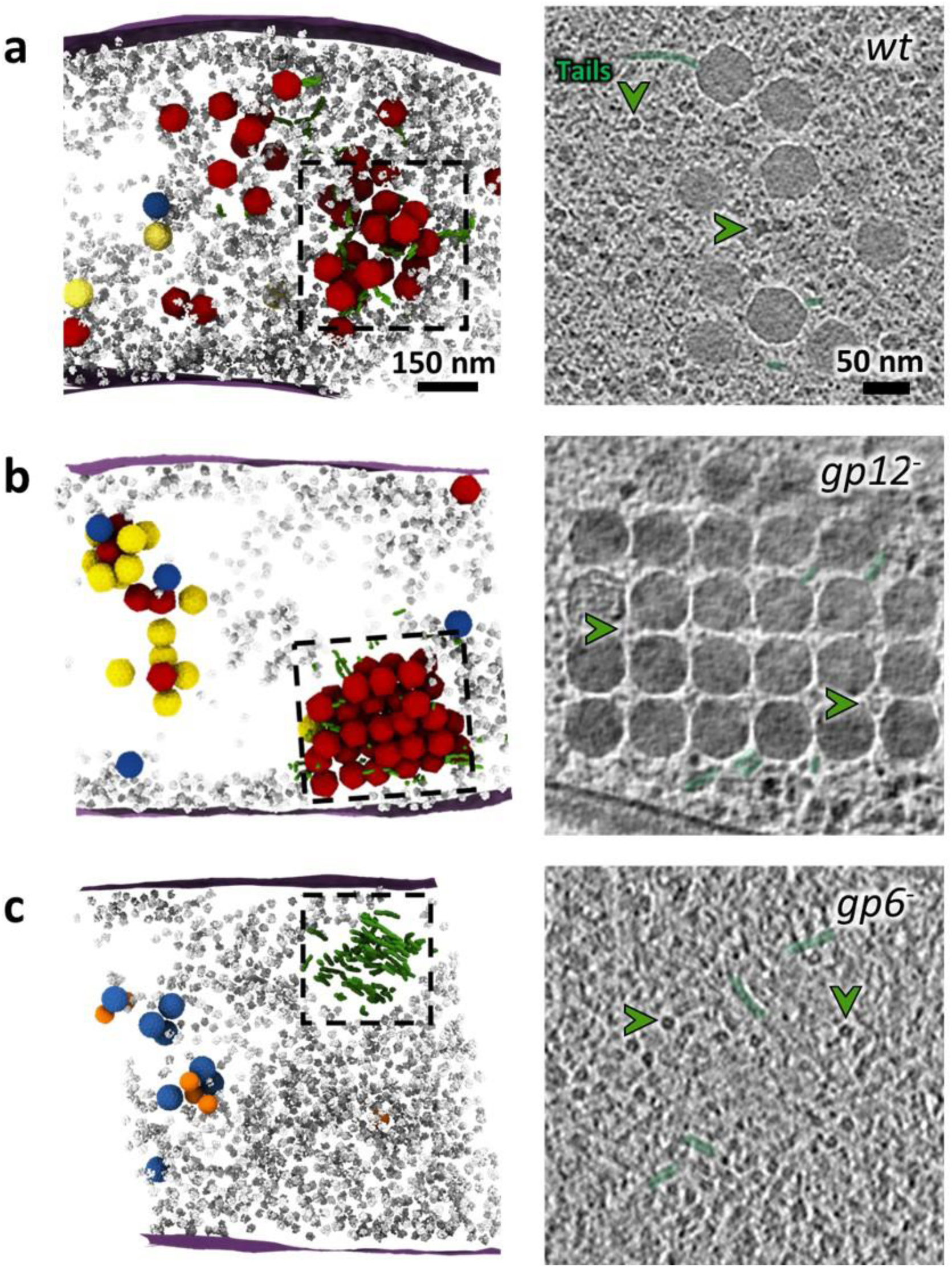
Clustering of viral particles outside the vDNA compartment and binding to pre-assembled tails after viral genome packaging. **a.** (*Left*) Segmentation rendering and back-plotting of a representative tomogram of *B. subtilis* infected by SPP1 wild type (*wt*) with a cluster of virions at the periphery of the vDNA compartment (Supplementary Movie 7). (*Right*) Detail of the cluster indicated by dashed lines on the left panel. **b.** (*Left*) Segmentation rendering and back-plotting of a representative tomogram of *B. subtilis* infected by SPP1lacO64gp12-(*gp12^-^*) showing a tightly packed warehouse at the periphery of the vDNA compartment (Supplementary Movie 8). (*Right*) Detail of the warehouse indicated by dashed lines on the left panel. **c.** (*Left*) Segmentation rendering and back-plotting of a representative tomogram of *B. subtilis* infected by SPP1*gp6^-^* (*gp6^-^*), defective in assembly of functional procapsids, displaying an aggregate of pre-assembled tails (Supplementary Movie 9). (*Right*) Detail of the tail aggregate indicated by dashed lines on the left panel. Segmented elements are the bacterial cell membrane (magenta), ribosomes (grey), procapsids I (blue), procapsids II (yellow), mature virions (red), tails (green), and small procapsid-like structures (orange). Some of the tails seen in cross-sections are labelled with a green arrowhead on the right panels.

These late events of viral assembly lead to the appearance of foci detected with a fluorescently labelled marker of DNA-filled capsids^7^ such as gp12-mCitrine^21^. By fluorescence microscopy, the foci are located in the vicinity of the vDNA compartment (Extended Data Fig. 4). In our tomograms, those regions appear as locally enriched in individual phage particles next to some pre-assembled tails in SPP1*wt* infections (Fig 5a; Extended Data Fig. 3f; Supplementary Movie 7). In contrast, during infection with a SPP1*gp12^-^* mutant strain, defective in production of the surface protein gp12, viral capsids are packed as large and ordered three-dimensional arrays termed warehouses (Fig. 5b; Extended Data Fig. 3g Supplementary Movie 8). We confirm that formation of phage warehouses is a result of capsid-capsid interactions depending on their surface properties and on the presence of the auxiliary protein gp12^21^. Indeed, particles lacking gp12 arrange as warehouses or clusters, which are significantly more compact than those containing gp12 (compare Fig. 5a and b).

Free tails are rare in infected cells when the whole assembly process of viral particles takes place such as in *wt* infections but can sometimes be observed at the periphery of the vDNA compartment (Fig. 5a). In contrast, clusters of tails are visible in the infected cell cytoplasm - again next to the vDNA compartment - when capsid assembly is arrested prior to tail-capsid association (Fig. 5c; Extended Data Fig. 3h; Supplementary Movie 9), as reported previously^7^. We conclude that phage tails are pre-assembled outside the vDNA compartment and binding to DNA-filled capsids after they depart from the vDNA compartment, leading to the formation of infectious particles in a spatially coordinated manner (Fig. 6).

**Fig. 6.**
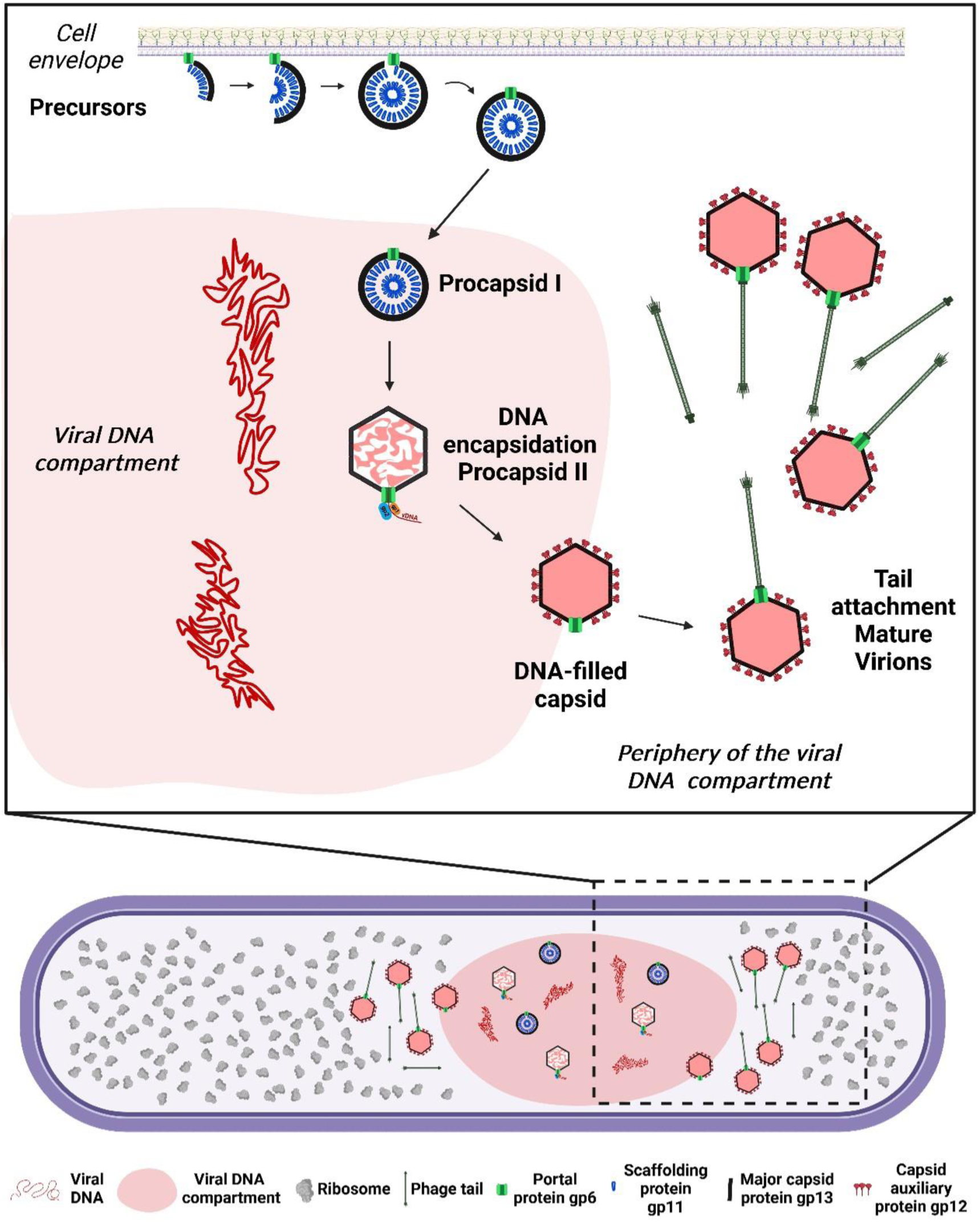
SPP1-induced spatial compartmentalization of viral particle assembly in the *B. subtilis* host cell. Model of SPP1 viral particle assembly progression inside a compartmentalized infected host bacterium. Following initiation of procapsid precursor assembly at the bacterial cell membrane, procapsids I relocate to the vDNA compartment where phage DNA replication and encapsidation take place. Initiation of viral DNA genome encapsidation triggers the structural transition from procapsid I to procapsid II that leads to release of scaffolding proteins and capsid expansion. Subsequently, DNA-filled capsids re-locate to the outside of the vDNA compartment and bind to pre-assembled tails there. Mature virions found at the periphery of the vDNA compartment can also cluster in warehouses within the bacterial cytoplasm prior to release by lysis of the bacteria host cell.

## Discussion

Our study reports the extensive reorganization of the bacterial cytoplasmic space induced upon siphovirus SPP1 infection and its role in orchestrating progeny virion assembly. Previously, we showed that a large vDNA compartment localizes asymmetrically in the *B. subtilis* infected cell and confines viral genome replication^7^. Using an optimized sample preparation workflow for cellular cryoET (Extended Data Fig. 1; see Online Methods), we here directly visualized under near-native conditions^9^ the architecture of this compartment that is delimited neither by a membrane nor by a protein cage (Fig. 1; Extended Data Fig. 3). This differs from giant jumbo phages whose vDNA is protected inside a proteinaceous lattice^4^. Nevertheless, the SPP1 vDNA compartment still excludes most of the cytoplasmic content such as ribosomes (Fig. 1b; Extended Data Fig. 4) and consequently protein translation reactions. Assembly of DNA-free procapsids I (Fig. 2) and tails (Fig. 5) also occurs outside the vDNA compartment and at different sites in the cell (Extended Data Fig. 3). Procapsids I, once assembled, relocate to the compartment (Extended Data Fig. 2, 9) where initiation of DNA packaging triggers their expansion to the procapsid II conformation (Fig. 4). DNA translocation into procapsids II takes place in a stepwise manner as unravelled in our tomograms by the succession of procapsid II particles partially filled with DNA (Fig. 4d-g; Extended Data Fig. 8). The resulting DNA-filled viral particles then depart and accumulate at adjacent positions, yet spatially separated from vDNA (Fig. 1b; Extended Data Fig. 9). Finally, DNA-filled capsids bind pre-assembled tails to form viral particles that can cluster and form paracrystalline arrays called warehouses depending on the presence of the auxiliary protein gp12 (Fig. 5). Altogether, we show that the intracellular assembly pathway of SPP1 follows a precise spatially regulated program in the compartmentalized bacterium (Fig. 6).

A major finding of our study is that open procapsid-like structures are observed and associated with the inner side of the bacterial cell membrane (Fig. 2; Extended Data Fig. 5) and that such localization depends strictly on the presence of the portal protein gp6 (Fig. 3; Extended Data Fig. 2). These unprecedented structures likely represent initial assembly stages of DNA-free procapsids I *in cellula*. We propose a model in which the portal protein bound to the inner side of the membrane promotes association of scaffolding proteins (gp11) that then serve to initiate unidirectional polymerization of the major capsid proteins (gp13). Next, growth of the icosahedral lattice is driven by co-assembly between the scaffolding and major capsid proteins until closure of the lattice to yield procapsid I (Fig. 2). In this assembly strategy, the precise curvature of the growing lattice observed in procapsid precursors (Fig. 2i) appears to be critical for formation of procapsids homogeneous in size. Interestingly, portals and/or procapsids of a few phages were previously shown to have a membrane-associated state (see ^4,22,23,24^). Overall, our proposed model illustrates how one single specialized DNA-translocating portal vertex can be assembled within an icosahedral lattice in a highly accurate and controlled manner (Fig. 6). Such association of phage portals with the bacterial cell membrane indeed provides a platform for procapsid assembly initiation and offers an effective way to prevent interference of additional portals with the scaffolding-major capsid polymerization reaction during the procapsid lattice growth phase in the cytoplasm.

Completion of procapsid I assembly marks its departure from the membrane to the vDNA compartment (Fig. 2d,h) where DNA packaging takes place (Fig. 4). Next, the DNA-filled capsids, bound to pre-assembled tails or not, are found outside of the compartment (Fig. 5). For each of these re-localization steps (Extended Data Fig. 9), our cryoET datasets provide no evidence for a dedicated transport system of viral particles that would be similar to the prokaryotic cytoskeleton utilized by jumbo phages in particular^4^. With this, changes in the properties of either the capsid surface or at the level of the membrane-less compartment in the course of the progressing infection represent the most likely drivers for the observed relocation events. Indeed, localization of procapsids I and procapsid I-like structures in the vDNA compartment is independent of the presence of the portal protein revealing that it relies on the procapsid lattice properties (Fig. 3e,f; Extended Data Fig. 9). Individual procapsids caught in the act of genome packaging inside the vDNA compartment were found to undergo expansion at the very early stages of DNA encapsidation and to leave the vDNA compartment as headful DNA-filled capsids (Fig. 4; Extended Data Fig. 8). We hypothesize that changes in the capsid lattice properties, resulting of its expansion combined with more subtle effects exerted by the tightly packaged vDNA on the capsid structure^14^, drive exit from the compartment. This is made possible after capsid detachment from vDNA concatemers upon genome packaging termination^7^. Binding of the auxiliary protein gp12 to the DNA-filled capsid hexamers prevents formation of tightly packed warehouse compartments of viral particles (Fig. 5) similar to those previously found in thin sections of cells infected with HSV-1^25^, podovirus P22^26^, and polyomavirus^27^. Collectively, these findings reveal that changes in the viral capsid surface properties during its assembly pathway is the main factor driving the spatial program of viral maturation in the compartmentalized infected cell.

Bacteriophages have been used to pioneer investigation of viral infection processes and to establish technology to probe the underlying molecular mechanisms over the past 6 decades (^28,29,30,31^). In combination with sophisticated biochemistry, research has provided detailed mechanistic and structural insights into bacteriophage assembly but lacked critical information on how they are modulated within the cellular environment. The present study expands on this extensive work and highlights the importance of the emerging field of cellular structural biology. Altogether, our new complementary findings provide a more comprehensive understanding of virus-host interactions within the bacterial cell context including in particular short-lived and transient stages of viral particle assembly inaccessible to purification for structural analysis *in vitro*.

## Supporting information

Supplementary Movie 1 | Representative tomogram of uninfected B. subtilis.

Supplementary Movie 2 | Representative tomogram of B. subtilis infected with SPP1 wt.

Supplementary Movie 3 | TIRF microscopy and time-lapse imaging of gp6-mCitrine in non-infected B. subtilis.

Supplementary Movie 4 | TIRF microscopy and time-lapse imaging of gp6-mCitrine in B. subtilis infected with SPP1lacO64gp6-.

Supplementary Movie 5 | Representative tomogram of B. subtilis infected with SPP1gp6-.

Supplementary Movie 6 | Representative tomogram of B. subtilis infected with SPP1lacO64gp2-.

Supplementary Movie 7 | Representative tomogram of mature virions warehouse in B. subtilis infected with SPP1 wt.

Supplementary Movie 8 | Representative tomogram of mature virions warehouse in B. subtilis infected with SPP1lacO64gp12-.

Supplementary Movie 9 | Representative tomogram of mature virions warehouse in B. subtilis infected with SPP1gp6-.

Supplementary Table 1 | Bacterial strains, viral strains, plasmids and oligonucleotides used in this work.

Supplementary Table 2 | Sample preparation and data collection parameters for electron cryo tomography.

## Online Methods

### Bacterial strains and growth conditions

Bacterial strains used in this work are listed in Supplementary Table 1. *B. subtilis* GSY10027 was constructed by transformation of *B. subtilis* GSY10004 with plasmid pAL27 linearized with ScaI. Double cross-over led to insertion of the gene coding for gp6-mCitrine under control of the inducible promoter *P_xyl_* at the *amyE locus*.

*B. subtilis* GSY10082 and GSY10087 were constructed in three steps. First, GSY10074 and GSY10085 were constructed by transformation of *B. subtilis* YB886 with plasmids pLG51 and pLG50, respectively, linearized with ScaI. Double cross-over led to insertion of genes coding for RpsB-mCFP and RplA-mCFP, respectively, under control of the inducible promoter *P_xyl_* at the *amyE locus*. Then, GSY10075 and GSY10086 were constructed by transduction of GSY10074 and GSY10085, respectively, with a lysate of SPP1 wild type (*wt*) amplified in strain GSY10004. The resulting strains encode LacIΔ11-mCherry under control of the constitutive promoter *P_pen_* at the *thrC locus*. Finally, GSY10082 and GSY10087 were constructed by transduction of GSY10075 and GSY10086, respectively, with a lysate of SPP1*wt* amplified in strain GSY10024. The resulting strains encode gp12-mCitrine under control of the inducible promoter *P_spac_* at the *sacA locus*. Expression from promoters *P_xyl_* and *P_spac_* was induced with 1% (w/v) xylose and 1 mM IPTG, respectively.

Transformation of competent *B. subtilis* cells was performed by a two-step starvation protocol using Spizizen medium for growth ^32^. Linearized plasmid (1 µg) was added to 500 µL competent cells for 30 minutes at 37°C, plated on LB plates supplemented with 5 µg/mL kanamycin for GSY10027 or 100 µg/mL spectinomycin for GSY10074 and GSY10085 and incubated overnight at 37°C. Transduction of *B. subtilis* cells was performed using previously published protocol^33^. To produce SPP1 lysates, donor strains GSY10004 and GSY10024 were grown overnight at 30°C in LB medium. Cultures were diluted 1:100 in 10 mL of LB medium and grown at 37°C. At an OD_600nm_ of 0.8, cultures were supplemented with 10 mM CaCl_2_, infected with SPP1*wt* at an input multiplicity of 5 pfu/cfu (*i.e*., plaque forming unit over colony forming unit) and incubated for 2 h at 30°C. Cell debris were sedimented by centrifugation at 4°C for 15 min at 8,000 *g* and the supernatant was stored at 4°C. Then, the SPP1 transducing lysates were used to infect 300 µL of a late-logarithmic-phase culture (OD_600nm_ of ∼1.6) of the receptor strain supplemented with 10 mM CaCl_2_ at an input multiplicity of 1. After 10 minutes at 37°C, the infected culture was mixed with 1.5 mL of prewarmed LB medium and further incubated with shaking for 10 minutes. Bacteria were sedimented and resuspended in 300 µL of prewarmed LB medium containing 40 μL of anti-SPP1 serum to inactivate free phages. Bacteria were incubated for 30 minutes, sedimented, and resuspended in 300 mL of prewarmed LB medium. Serial dilutions of the culture were plated in solid medium supplemented with appropriate antibiotics and incubated at 37°C overnight. Antibiotic selection was 12.5 µg/mL lincomycin + 0.5 µg/mL erythromycin (*mls* resistance) and 5 µg/mL kanamycin (*kan* resistance).

### Plasmid construction

Plasmids and oligonucleotides used in the present study are listed in Supplementary Table 1 with references. SPP1 gene *6*, coding for the portal protein, was amplified by PCR from SPP1*lacO64* DNA with primers 55 and 56. The PCR product digested with ClaI-XhoI was ligated to pAL21 cut with the same restriction enzymes to generate plasmid pAL27.

Genes *rplA*, coding for ribosomal protein L1, and *rpsB*, coding for ribosomal protein S2, were amplified by PCR from *B. subtilis* strain YB886 DNA with primers 20 and 28, 22 and 29, respectively. PCR products digested with KpnI-EcoRI for *rplA* or with KpnI-XhoI for *rpsB* were ligated to pAL29 cut with the same enzymes, generating pLG50 and pLG51, respectively. These constructs encode L1 and S2, respectively, fused to CFP at their carboxyl terminus. The *cfp* synthetic gene was codon-optimized for *B subtilis*. Fusion genes were expressed under the control of a xylose-inducible promoter.

Plasmid constructions were transformed into *Escherichia coli* DH5α and selected on LB plates supplemented with 100 µg/mL ampicillin. Clones were checked by DNA sequencing.

### Construction of phage strains

Phage strains used in this work are listed in Supplementary Table 1. Phage SPP1*delX110lacO64sus115* (abbreviated SPP1*lacO64gp6^-^*) was generated by co-infecting the permissive *B. subtilis* strain HA101B with phages SPP1*delX110lacO64* (abbreviated SPP1*lacO64*) and SPP1*sus115* (abbreviated SPP1*gp6^-^*) at an input multiplicity of 10 pfu/cfu for each phage strain. Co-infection was carried out in LB medium supplemented with 10 mM CaCl_2_ at 37°C for 2 hours. The cell lysate was centrifuged at 12,000 *g* at 4°C for 20 minutes and the supernatant was titrated using bacterial strain HA101B. Ninety-six isolated phage plaques were collected and resuspended separately in 200 µL of TBT buffer (100 mM Tris-Cl pH 7.5, 100 mM NaCl, 10 mM MgCl_2_) in a 96-well plate. Phage clones were spotted in the non-permissive strain YB886, in the permissive strain HA101B and in the complementing strain YB886 (pSPW7) (Supplementary Table 1). Phages carrying mutation *sus115* were identified by not multiplying in strain YB886. These clones were then screened by PCR to identify those carrying insertion *lacO64* (primers 3185 and 3682; Supplementary Table 1) and deletion *delX* combined with insertion *lacO64* (primers 2677 and 3383; Supplementary Table 1)^33^. Selected clones were re-titrated to obtain single plaques of pure clones, their genotype confirmed as above and amplified in bacterial strain HA101B.

### Fluorescence microscopy

Fluorescence microscopy of GSY10082 and GSY10087 strains was performed as previously described^7^. For Total Internal Reflection Fluorescence Microscopy (TIRFM) experiments, samples of bacterial cells were cultivated and infected with an input multiplicity of 2 pfu/cfu. After ∼23 minutes of incubation at 37°C with orbital shaking, cells were spotted on a 1% agarose pad (thickness of 2 mm) diluted in CH medium. Initial single exposure images were obtained with conventional epifluorescence images using 100 ms-long exposures at 488 nm and 561 nm. Then, time-lapse images were collected with TIRF using 100 ms-long exposures at 488 nm and 561 nm with an image taken every 2 seconds for 1 or 2 minutes. Final control images illuminated by epifluorescence were collected from a single exposure of 100 ms at 488 nm. TIRFM imaging was performed using a Nikon Ti-Eclipse microscope with a 100X oil objective (NA: 1.3; WD: 0.2 mm; CFI Plan Fluor - Phase - Nikon) and an EMCCD camera (Andor). Snapshots were extracted from time-lapse movies.

### Electron cryo tomography

#### Sample preparation

*B. subtilis* strain GSY10024 was cultivated in LB medium supplemented with 1 mM IPTG at 37°C. At OD_600nm_=0.8 (∼10^8^ cfu/mL), the culture was supplemented with 10 mM CaCl_2_ and cells were infected by SPP1 phages at 37°C for ∼15 minutes (early time point) or ∼25 minutes (late time point) at an input multiplicity of 5 pfu/cfu. Cells were then centrifuged at 16,200 *g* for 1 minute at room temperature and pellets were resuspended in MIII medium^34^ in 1/10 of the culture initial volume. Non-infected cells were treated the same way as a control. For electron cryo tomography (cryoET), 4 µL of infected or non-infected bacteria were added to electron microscopy (EM) grids (Quantifoil Micro Tools GmbH; Cu R2/2) before blotting for 20 s on one side and plunge freezing using a Leica EM GP plunger (Leica Biosystems Nussloch GmbH) (Supplementary Table 2).

Grids were clipped using C-clips and autoGrids in the AutoGrid assembly workstation (ThermoFisher Scientific). Lamellae were prepared in an Aquilos 2 (ThermoFisher Scientific) equipped with a gallium focused ion beam (FIB in short), used to make thin sections of cells called lamellae by ablating cellular material around the area of interest. For this, sample preparation, milling and polishing steps were done automatically using Maps and AutoTEM software (ThermoFisher Scientific) with a milling angle target of 8 degrees (2 degrees of tolerance) and a final lamella thickness set to 120 nm. The grids were retrieved after polishing of the lamellae and stored for a minimal amount of time in liquid nitrogen prior to data collection.

#### Data acquisition

Grids with milled lamellae were transferred for tilt-series acquisition to a Titan Krios G3 operated at 300 kV (ThermoFisher Scientific), equipped with a K3 direct electron detector and a post-column BioQuantum energy filter (Gatan). Grids were screened and data acquired using SerialEM^35^. Grid scans were acquired to locate the lamellae. Tilt-series were collected using a dose-symmetric tilt scheme^36^ from 60 to −60 degrees specimen tilt range with a 3 degree increment step starting with 7-10 degree of pre-tilt (according to the specific milling angle of each lamella). Frames were acquired at 15 e-/pixel/s in electron counting mode with a pixel size of 3.36, 2.15 or 1.37 Å and aligned using SerialEM implementation (see Supplementary Table 2). A 70 μm objective aperture and a 20 eV energy slit were inserted during data acquisition. A nominal defocus range between 4 and 5 µm was used.

#### Tomogram reconstruction

Reconstruction of tomograms from tilt-series was done using IMOD version 4.12.40^37^ or Aretomo^38^ when no fiducial-like feature were visible on the projection images. For visualization purposes, we typically used a binning of 4 and deconvolution using the standard parameters of ISONET without missing wedge compensation^39^.

#### Sub-volume averaging of procapsid I

572 procapsids I were manually picked using IMOD^37^ from tomograms binned by 4 (n = 30; pixel size = 8.6 Å) and separated in two halves for independent processing done in PEET 1.14.1^40,41^. For each dataset, the initial reference was the *in vitro* determined map of Procapsid I (EMD-4717)^14^ lowpass filtered to 55 Å. This reference was used for averaging with particles having randomized orientations using custom scripts based on TEMPy^42^ with a box size of 80*80*80 voxels. The MCP of this average was segmented using the segger tool^43^ in Chimera^44^, to obtain a mask for subsequent alignment. The whole capsid average volume using this mask converged to (stabilised on) an icosahedral structure. Alignment was then focused on the 12 icosahedral vertices using custom scripts based on TEMPy, resulting in 6864 particles extracted with a box size of 80*80*80 voxels. Further iterative alignment steps were done until convergence. The final average volume reached a resolution estimated to 26.5 Å by Fourier Shell Correlation using PEET calcUnbiasedFSC.

#### Volume visualisation

surface rendering of volumes was performed using ChimeraX. Segmentation of bacterial cell membrane was done using drawing tools in 3dmod. Average volumes of procapsids I, procapids II, mature virions, and ribosomes were placed back within each tomogram using coordinates and orientations obtained by sub-volume averaging with PEET 1.14.1^40,41^ using custom scripts based on TEMPy^42^ as described above. Here, sub-volume averaging was performed using the following references: procapsid I (EMD-4717), procapsid II (EMD-10002), mature virions (EMD-4716), and ribosomes (EMD-5787). Template matching for ribosomes was done by converting heavily filtered and thresholded volumes to model point coordinates and aligning these to a filtered reference. After cleaning by cross-correlation and removal of duplicate particles, the resulting sub-volumes were manually curated from false positive and plotted back in the original tomogram coordinate system.

### Statistical analysis

Statistical analysis was performed using RStudio software. All data is represented as a boxplot: the full line represents the median and each point corresponds to one tomogram. Shapiro and Bartlett’s tests were used to evaluate normality and variance homogeneity. As the data distribution was not normal and/or variances were not homogenous (p-value<0.05), non-parametric tests were used for statistical analysis. Statistical significance for the ratio of the number of capsids inside the vDNA compartment to the total number of capsids was evaluated using either a Kruskal-Wallis test (non-parametric one-way ANOVA test) or a Wilcoxon test (non-parametric t-Student test). Statistical significance is assessed by a p-value < 0.05.

### Data visualization

Figures were prepared with Adobe Illustrator, Biorender and Chimera^44^.

## Data Availability

Average map of the procapsid I included in this paper has been deposited in the Electron Microscopy Data Bank (EMDB) with the following accession code: EMD-54016

## Acknowledgements

We acknowledge funding for this collaborative project by ANR SelectVir (ANR-23-CE12-0019-01) to P.T and E.Q. In the framework of this project, S.C.D. benefited from a travel grant from the Leibniz Institute of Virology (LIV) and a Mobility fund from the Deutsch-Französiche Hochschule IB-ID doctoral college to visit I2BC. E.Q. was supported by a Klaus Tschira Boost Fund in 2021 and received funding from the ATIP-Avenir programme in 2022.

Part of this work was performed at the CryoEM multi-user Facility at CSSB, headed by K.G. and supported by the UHH and DFG (grants INST 152/772-1, 774-1, 775-1 and 777-1). We also acknowledge excellent support by Carolin Seuring, Cornelia Cazey, and Ulrike Laugks, for sample preparation and data acquisition, and Wolfgang Lugmayr for supporting workflows for cryoEM/ET data processing on the Maxwell compute cluster. Grids were prepared using the Synchrotron SOLEIL facilities.

We are indebted to Cyrille Billaudeau and Rut Carballido-Lopez (Micalis, INRAE, Jouy-en-Josas) to make generously available the Nikon epifluorescence microscope equipped with a TIRF system, for their continuous support for these experiments, and for discussions on results interpretation.

## Author contributions

SCD, AL, KG, PT and EQ conceived the project and obtained funding. SCD, AL, PT and EQ designed the experiments. SCD, AL, PL and EQ optimized sample preparation on grids. Cloning, virus production, protein expression and complementation assay was done by AL, LG and PT. SCD performed sample preparation and data collection for cryoET with supervision from EQ. AL carried out the TIRF experiments and analysis. SCD, VP and EQ did the processing of cryoET datasets and sub-volume averaging. CM was responsible for the statistical analysis. SCD, AL, PT and EQ prepared the manuscript with input from all co-authors.

## Competing interest declaration

The authors declare that they have no known competing interests.

## Extended Data

**Extended Data Fig. 1.**
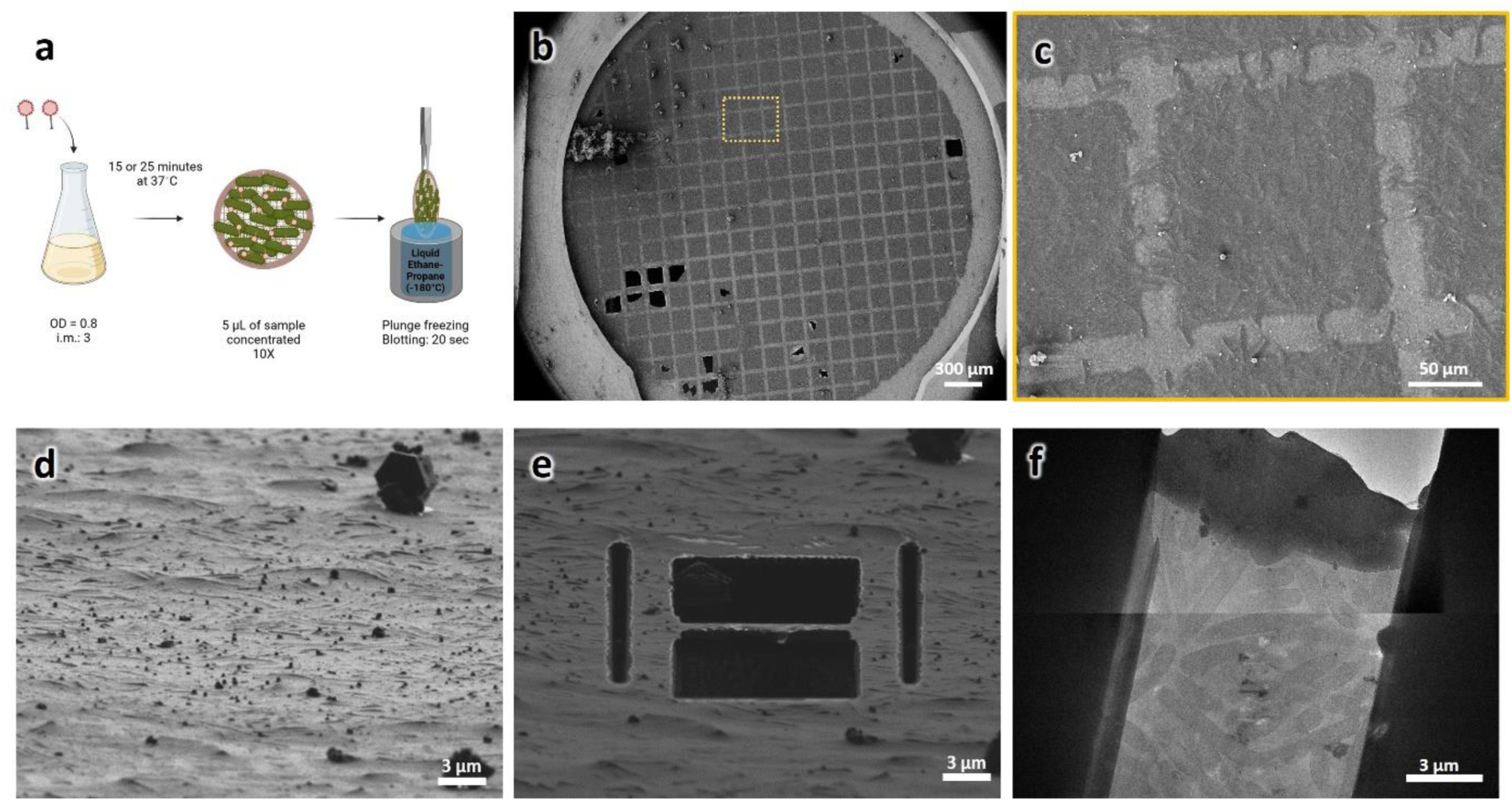
Sample preparation workflow developed for studying SPP1-infected bacteria by electron cryo tomography. **a.** Schematic representation of the optimized protocol used for sample preparation for cryoET. Following *B. subtilis* infection by SPP1, cells concentrated 10-fold are applied on grids and plunge-frozen in cryogen cooled down to liquid nitrogen temperature. OD, optical density; i.m., input multiplicity. **b.** Overview of a sample plunge-frozen on grid observed under cryo conditions in a scanning electron microscope (SEM) equipped with a focused ion beam (FIB), a cryo-FIB-SEM. The yellow rectangle on the SEM view indicates the region enlarged in (**c**) showing a grid square with a continuous layer of *B. subtilis* cells on top. **d,e.** FIB view of the targeted grid area before (**d**) and after FIB-milling (**e**) using the cryo-FIB-SEM to remove material at the top and bottom of the area of interest to make lamellae that are ∼100 nm thin. **f.** Overview of a representative lamella containing cross-sections of several *B. subtilis* cells observed after transfer to a Titan Krios for cryoET data collection. Scale bars are indicated.

**Extended Data Fig. 2.**
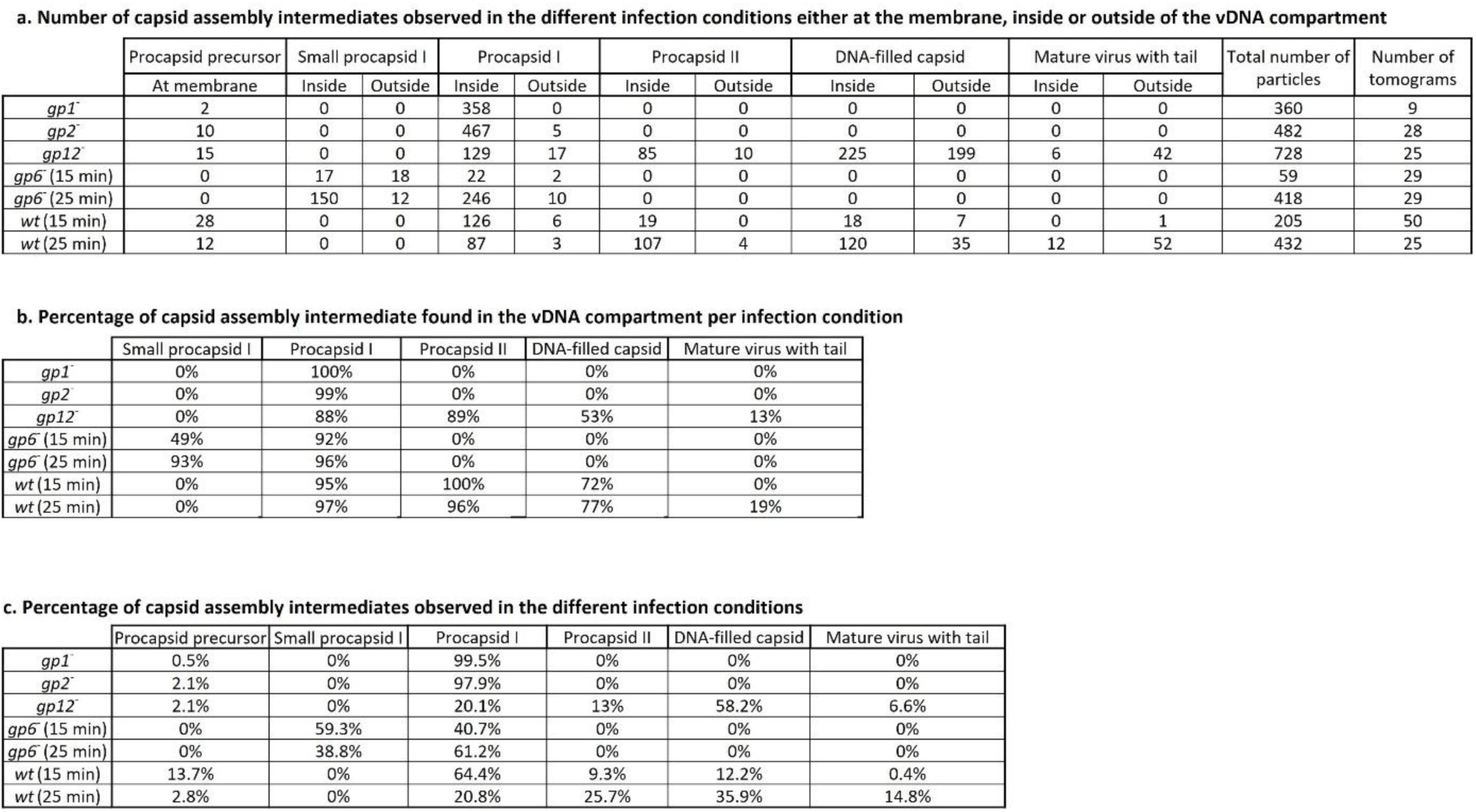
Quantification of capsid structures observed in wild type and mutant SPP1 infection conditions. **a.** Number of capsid assembly intermediates and their location into the compartmentalized host bacterial cell observed in the tomograms under the different conditions of infection. The localisation inside the infected host bacterial cell have been divided as either “at the cell membrane” for precursors found at the membrane or “inside” and “outside” the vDNA compartment for the other stages of assembly observed. **b.** Percentage of the capsid assembly intermediates found specifically in the vDNA compartment in the different infection conditions analysed. **c.** Percentage of capsid assembly intermediates observed in the different infection conditions compared to the total number of events.

**Extended Data Fig. 3.**
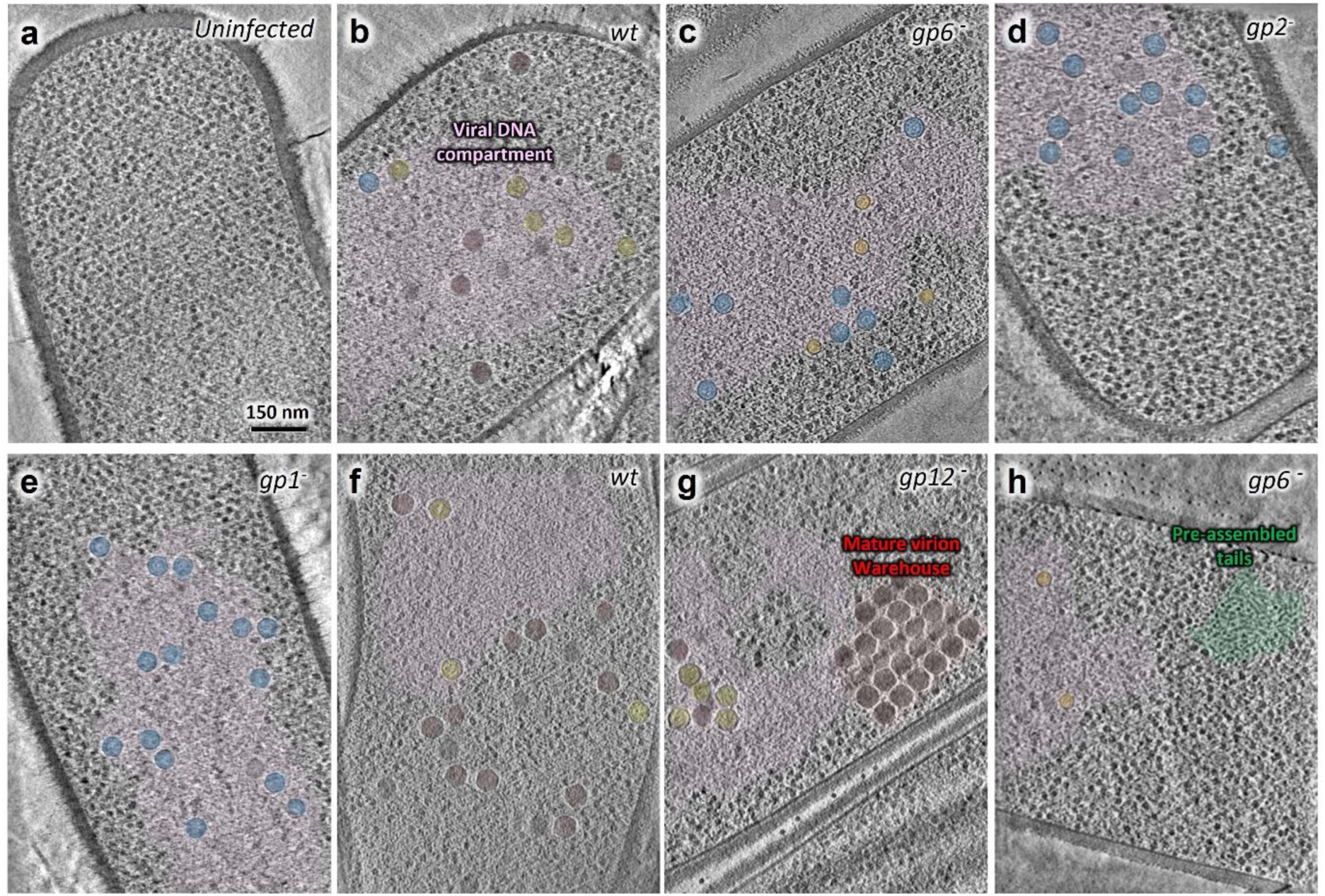
Overview of the vDNA compartment and viral particle structures observed in bacteria infected by wild type and mutant SPP1 phages. a. Slice from a representative tomogram of uninfected *B. subtilis* cell (shown in Fig. 1a; Supplementary Movie 1). b. Slice from a representative tomogram of *B. subtilis* infected by SPP1 wild type (*wt*; shown in Fig. 1b; Supplementary Movie 2) with the vDNA compartment visible (pink) containing different assembly intermediates: procapsids I (blue), procapsids II (yellow), and DNA-filled capsids (red). c. Slice from a representative tomogram of *B. subtilis* infected by SPP1*gp6^-^*(*gp6^-^*; shown in Fig. 3c; Supplementary Movie 5) with the vDNA compartment visible (pink) containing procapsids I (blue) and small procapsid-like structures (orange). d. Slice from a representative tomogram of *B. subtilis* infected by SPP1lacO64*gp2^-^* (*gp2^-^*; shown in Fig. 4a; Supplementary Movie 6) with the vDNA compartment visible (pink) containing exclusively procapsids I (blue). e. Slice from a representative tomogram of *B. subtilis* infected by SPP1*gp1^-^*(*gp1^-^*) with the vDNA compartment visible (pink) containing exclusively procapsids I (blue). f. Slice from a representative tomogram of *B. subtilis* infected by SPP1 wild type (*wt*; shown in Fig. 5a; Supplementary Movie 7) with the vDNA compartment visible (pink) and a diffuse warehouse at its periphery containing mature virions (red). g. Slice from a representative tomogram of *B. subtilis* infected by SPP1*lacO64gp12^-^* (*gp12^-^*; shown in 5b; Supplementary Movie 8) with the vDNA compartment visible (pink) and a packed warehouse at its periphery containing mature virions (red). h. Slice from a representative tomogram of *B. subtilis* infected by SPP1*gp6^-^*(*gp6^-^*; shown in Fig. 5c; Supplementary Movie 9) with the vDNA compartment visible (pink) as well as preassembled tails (green) clustered at its periphery.

**Extended Data Fig. 4.**
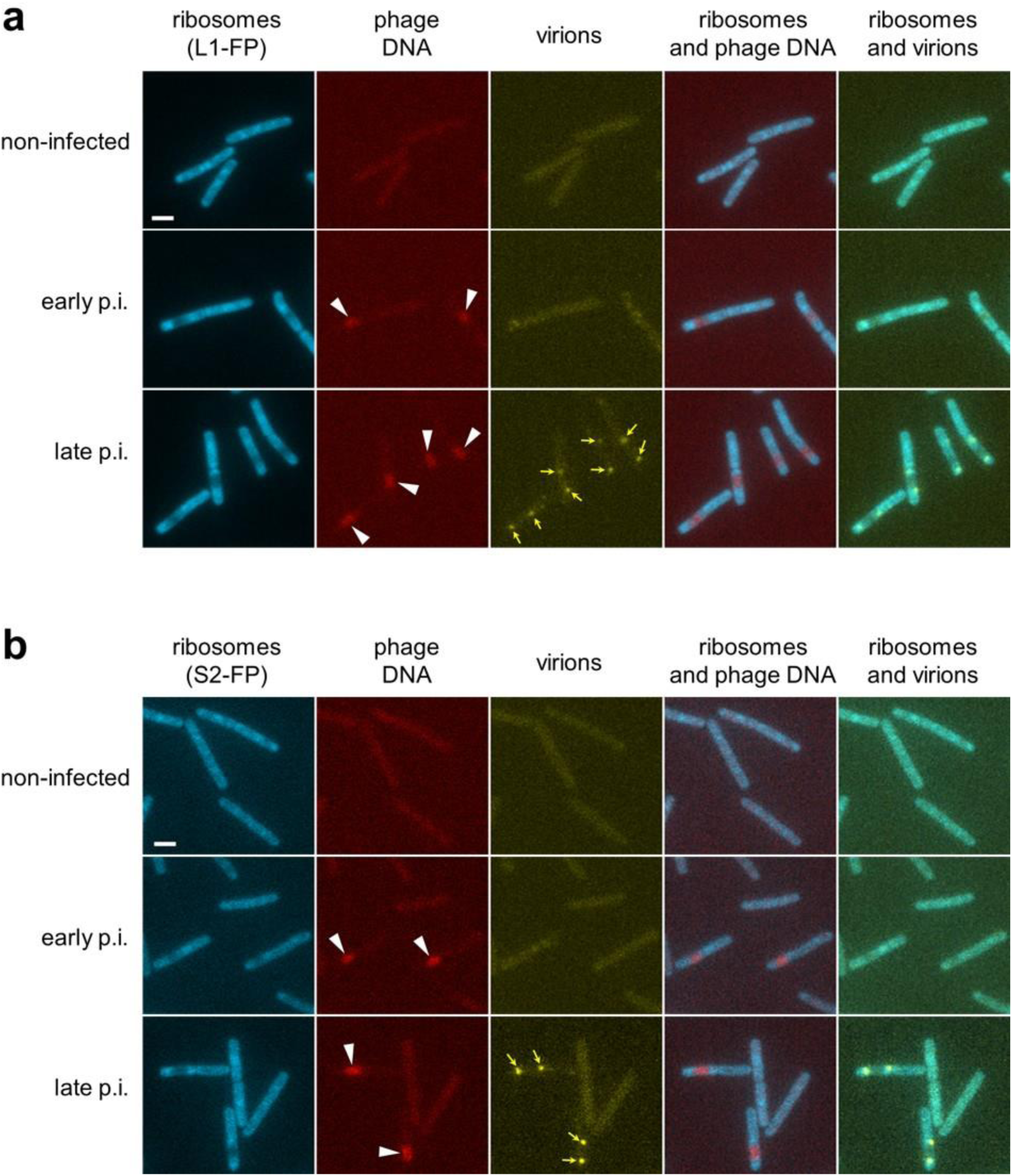
Localization of ribosomes, SPP1 DNA and virions inside the reorganized bacterial cells. Ribosomes are viewed by imaging ribosomal proteins L1 (or RplA in *B. subtilis*) (**a**) and S2 (or RpsB in *B. subtilis*) (**b**) fused to CFP (see Online Methods). Bacteria infected by SPP1*lacO64gp12^-^*were imaged at 30 min p.i. (early p.i.) and 55 min p.i. (late p.i.). Phage DNA is visualized with LacI-mCherry to define the vDNA compartment highlighted by white arrowheads in the RFP channel images. Virions are visualized with gp12-mCitrine and their warehouses highlighted by yellow arrows in the YFP channel images. Scale bars represent 2 µm.

**Extended Data Fig. 5.**
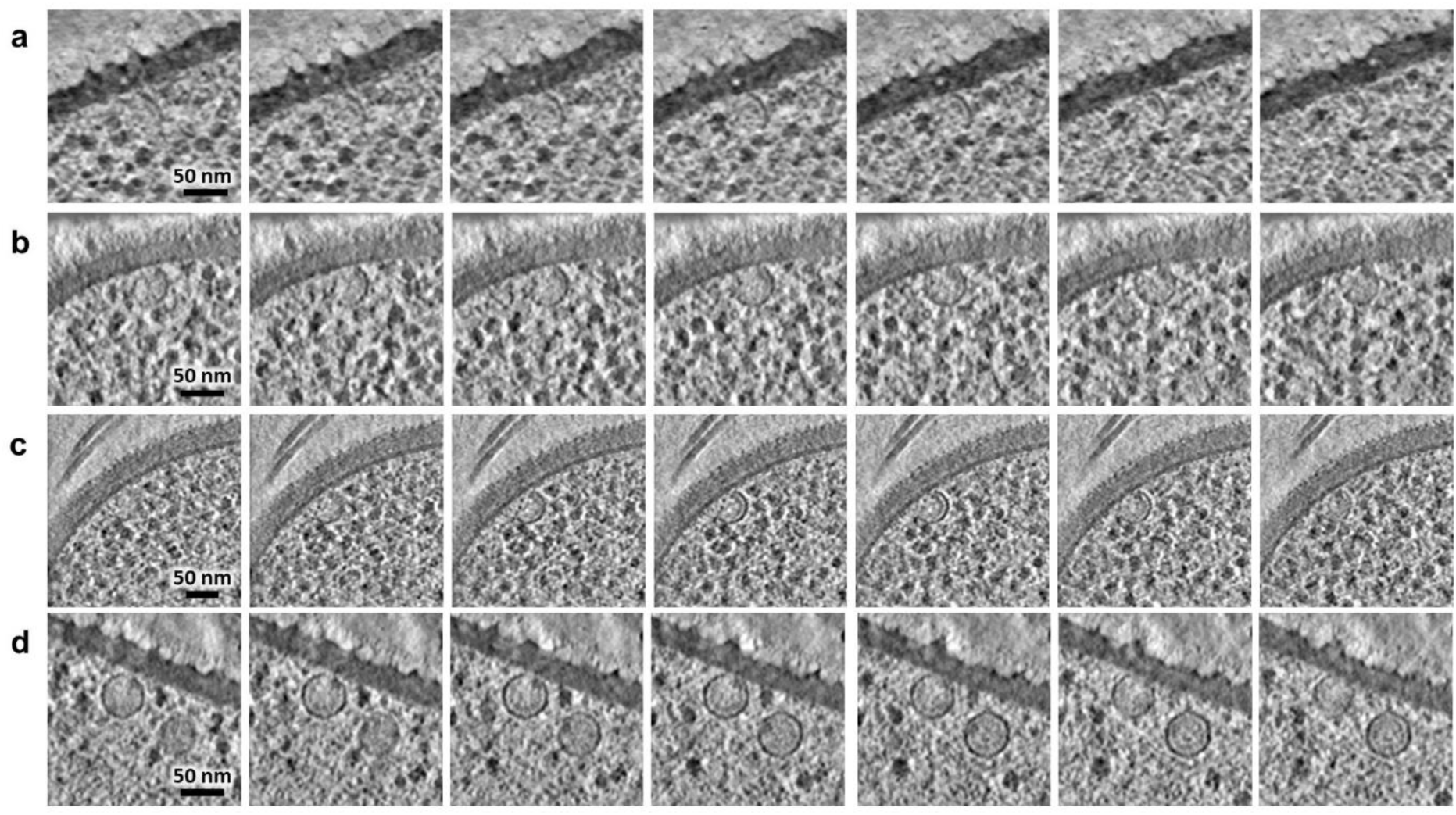
Procapsid assembly precursors. Gallery of slices through representative tomograms of *B. subtilis* infected by SPP1 wild type. Intermediates at different stages of procapsid assembly are showed at the bacterial cell membrane from an initial stage (**a**) to a complete procapsid I-like particle associated to the inner side of the bacterial cell membrane or detached (**d**). Each precursor of assembly intermediates is depicted here at different z heights while only a central slice is shown in Fig. 2. Z step: 2.6 nm for (**a,b**) and 3.4 nm for (**c,d**).

**Extended Data Fig. 6.**
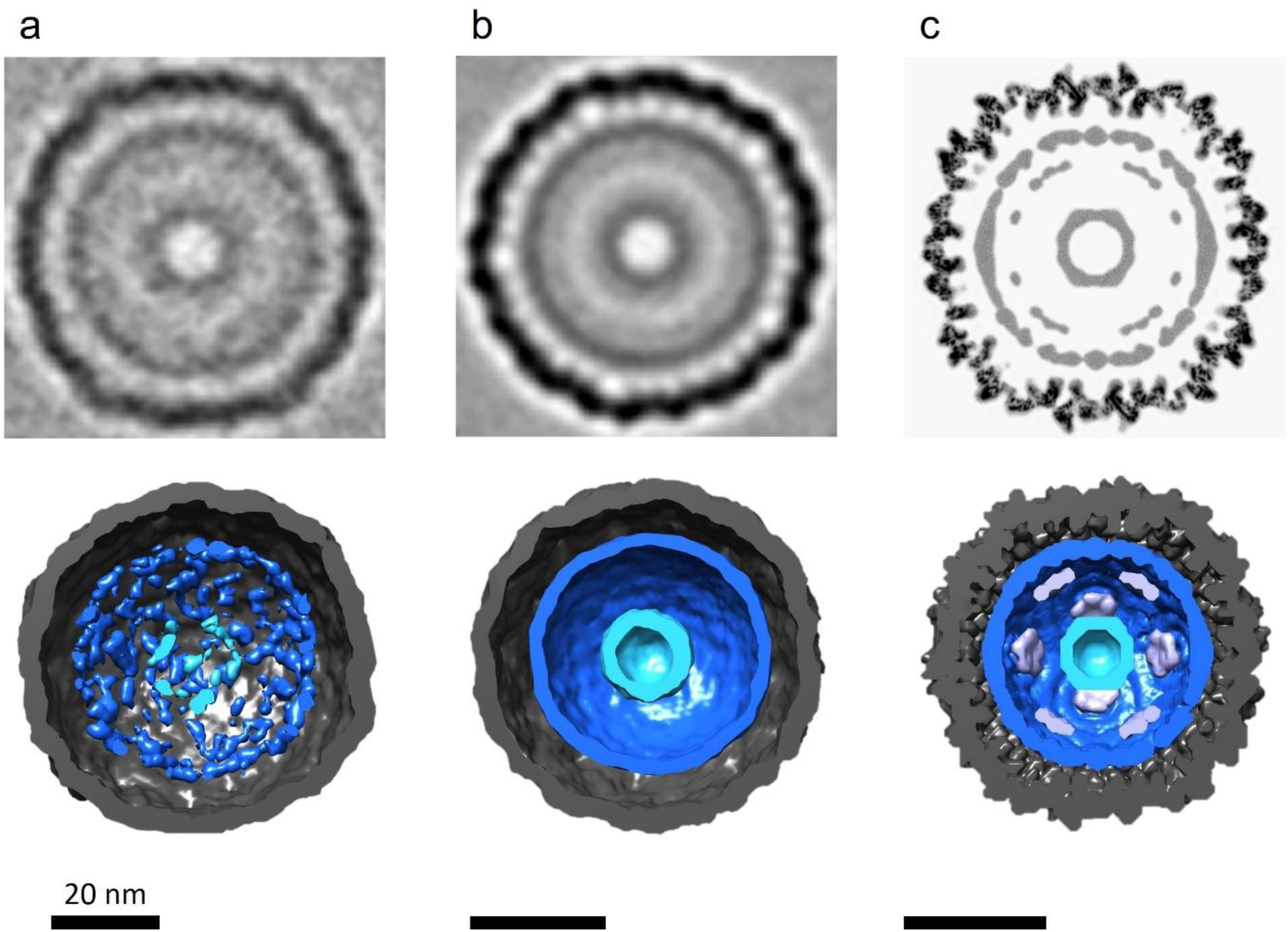
Sub-volume averaging and comparison of procapsid I structures found *in cellula* and *in vitro*. Gallery of slices through averages of procapsids I from a dataset in cellula of SPP1 wild type, SPP1lacO64gp1- and SPP1lacO64gp2-infection, accompanied by their respective volume rendering (bottom), compared to a cryoEM structure of in vitro purified particles published previously (EMD-4717). a. Average using one model point per capsid located at its centre (n = 572). b. Average after applying icosahedral symmetry using model points located at the centre of the capsid’s pentons (n = 6864). c. Published EM map from the previous cryoEM single-particle analysis of in vitro purified particles (Ignatiou et al 2019). Densities of features of interest are displayed with the inner shell in light blue and outer shell in dark blue, corresponding to the scaffolding protein, as well as intermediate densities in purple and the capsid in black. Scale bars are indicated.

**Extended Data Fig. 7.**
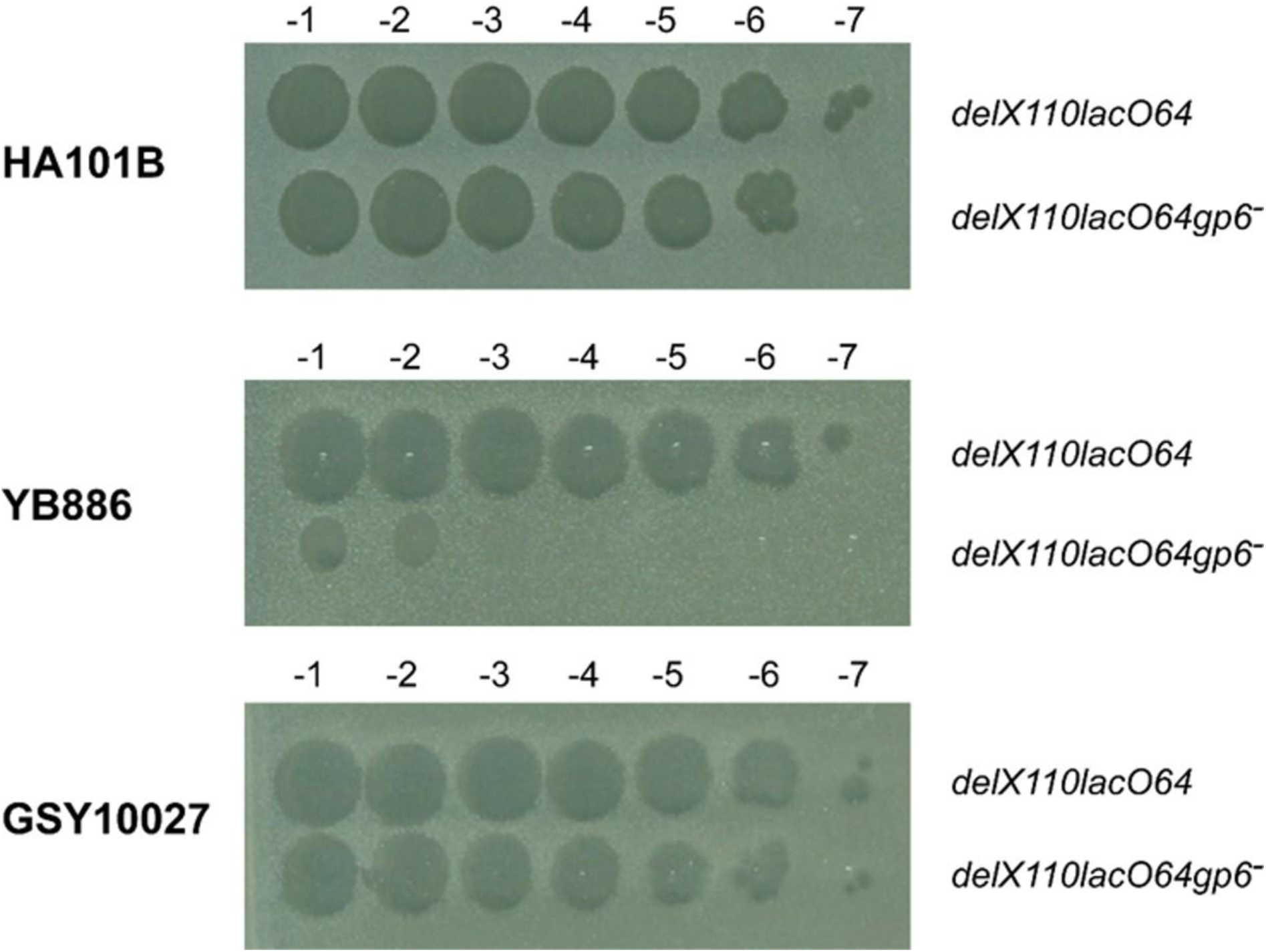
Trans-complementation of SPP1*gp6^-^* with gp6-mCitrine *in vivo*. Infection of *B. subtilis* permissive strain HA101B, non-permissive strain YB886, and strain GSY10027 by the suppressor-sensitive mutant SPP1*lacO64gp6^-^* and by the control phage SPP1*lacO64* (*wt*). GSY10027 is an YB886-derived strain that encodes gp6-mCitrine (see Online Methods). A 2 µL spot of each phage serial dilution was deposited on top of the indicator bacteria lawns. Phage stocks were initially diluted to 10^8^ pfu.mL^-1^ for serial dilutions from 10^-1^ to 10^-7^ as indicated on top.

**Extended Data Fig. 8.**
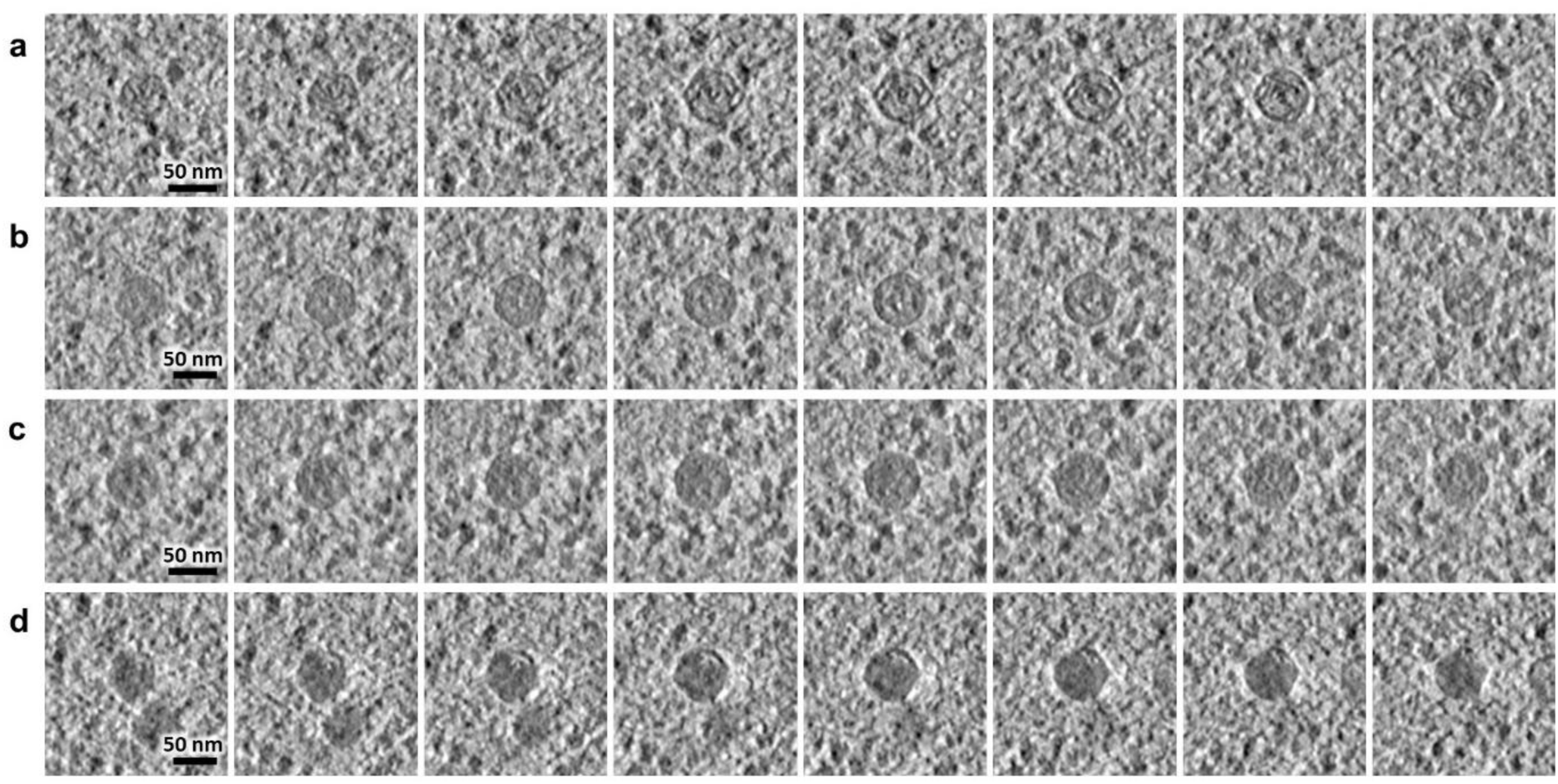
Procapsids II at different stages of viral genome encapsidation. Gallery of slices through representative tomograms of *B. subtilis* infected by SPP1 wild type. The gallery displays several procapsids II from an initial stage of genome packaging with a low amount of DNA observed inside the procapsid (**a**) to an almost filled capsid (**d**) and intermediates (**b** and **c**). Individual procapsids II are depicted here at different z heights to highlight how the viral genome is arranged on the inside during the process of encapsidation. Z step is 3.4 nm and scale bars are indicated.

**Extended Data Fig. 9.**
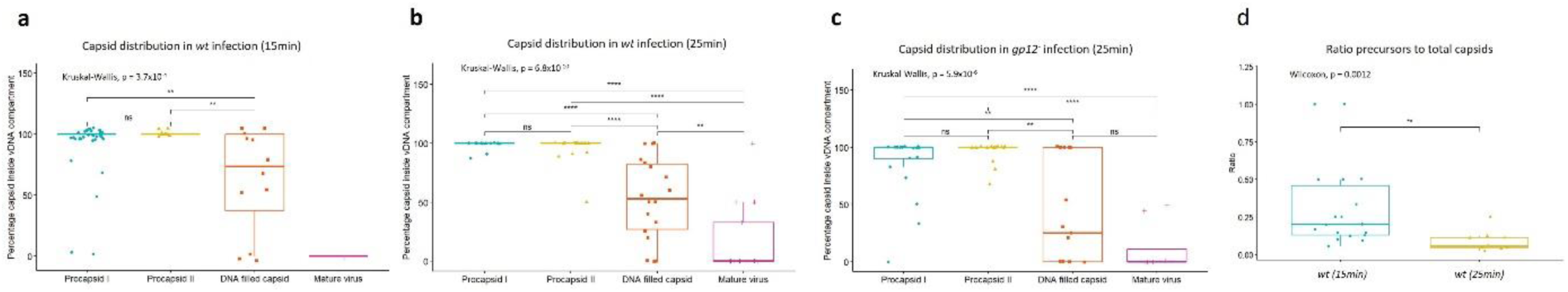
Statistical analyses on the timing and localization of the viral assembly intermediates in SPP1-infected cells. **a-c**. Percentage of the different assembly intermediates (procapsids I, procapsids II, DNA-filled capsids and mature virions) found inside the vDNA compartment at 15 min p.i. (**a**) and 25 min p.i. (**b**) in SPP1*wt* infection or at 25 min p.i. for SPP1*gp12-* infection (**c**). Regarding the distribution, the boundaries of the vDNA compartment were arbitrarily defined by the exclusion of the ribosomes (as exemplified in Fig. 1; Extended Data Fig. 3). **d**. Ratio of precursors compared to the total number of capsids (procapsids I, procapsids II, DNA-filled capsids and mature virions) is plotted for each tomogram. Precursors are considered as incomplete procapsid structures found at the bacterial cell membrane (as in Fig. 2; Extended Data Fig. 5). Statistically-relevant differences were determined using either a Kruskal-Wallis test (non-parametric one-way ANOVA test) or a Wilcoxon test (non-parametric t-Student test). Statistical significance is indicated by star symbols and assessed by a *p*-value below 0.05 (*), below 0.01 (**) or below 0.001 (****); ns, not significant. Each dot in the box plot corresponds to one tomogram analysed: n = 50 (SPP1*wt* at 15 min p.i.), n=25 (SPP1*wt* at 25 min p.i.), n=25 (SPP1*lacO64gp12*^-^ at 25min p.i.). The horizontal line represents the average for each condition.

## Supplementary information

### Supplementary Tables

**Supplementary Table 1 | Bacterial strains, viral strains, plasmids and oligonucleotides used in this work.** List of biological resources including bacterial strains, bacteriophage strains, plasmids and sequence of oligonucleotides employed in this work together with description of the source and short names used in the text.

**Supplementary Table 2 | Sample preparation and data collection parameters for electron cryo tomography.** List of samples with the number of tilt-series collected in parenthesis. Then information for sample preparation on grid by vitrification with the LEICA EM GP is given together with the parameters used for data collection for all datasets reported in this study.

### Supplementary Movies

**Supplementary Movie 1 | Representative tomogram of uninfected *B. subtilis***. The tomogram displays the typical cellular organization observed with the cytoplasm full of ribosomes except for a small area that contains the bacterial DNA. Segmentation rendering of this tomogram is illustrated in Fig. 1a and an original slice is showed in Extended Data Fig. 3a.

**Supplementary Movie 2 | Representative tomogram of *B. subtilis* infected with SPP1 *wt***. The tomogram exhibits the typical host cell re-organization observed upon SPP1 *wt* infection with a large area containing viral DNA and some capsid assembly intermediates with a clear exclusion of ribosomes. Segmentation rendering of this tomogram is illustrated in Fig. 1b and an original slice is showed in Extended Data Fig. 3b with colour overlay.

**Supplementary Movie 3**| TIRF microscopy and time-lapse imaging of gp6-mCitrine in non-infected ***B. subtilis***. *B. subtilis* cells producing gp6-mCitrine were imaged under TIRF illumination for 1 min with an acquisition every 2 s. Gp6-mCitrine small foci found in the perimembrane region are highly mobile. Snapshots of the movie are shown on the right panels of Fig. 3a.

**Supplementary Movie 4 | TIRF microscopy and time-lapse imaging of gp6-mCitrine in *B. subtilis* infected with SPP1*lacO64gp6^-^*.** *B. subtilis* cells producing gp6-mCitrine were infected with SPP1*lacO64gp6^-^*for 23 min at 37°C and imaged under TIRF illumination for 1 min with an acquisition every 2 s. Most gp6-mCitrine gets recruited to immobile intense foci localized outside the viral DNA compartment. Snapshots of the movie are shown on the right panels of Fig. 3b.

**Supplementary Movie 5 | Representative tomogram of *B. subtilis* infected with SPP1*gp6^-^***. The tomogram exhibits the typical host cell re-organization observed upon infection with phage SPP1*gp6*^-^ that is defective in production of the portal protein. Procapsid I-like structures and small procapsids I are found in the viral DNA compartment while aberrant capsid-like structures are visible at its periphery. Segmentation rendering of this tomogram is illustrated in Fig. 3c and an original slice is showed in Extended Data Fig. 3c with colour overlay.

**Supplementary Movie 6 | Representative tomogram of *B. subtilis* infected with SPP1*lacO64gp2^-^***. The tomogram exhibits the typical host cell re-organization observed upon infection with SPP1*lacO64gp2*^-^ that is defective in viral DNA packaging. Only procapsids I are found in the viral DNA compartment while one procapsid precursor is visible at the cell membrane. Segmentation rendering of this tomogram is illustrated in Fig. 4a and an original slice is showed in Extended Data Fig. 3d with colour overlay.

**Supplementary Movie 7** | Representative tomogram of mature virions warehouse in *B. subtilis* infected with SPP1 *wt*. The tomogram exhibits the typical host cell re-organization observed upon infection with SPP1 *wt*. Mature virions are found as clusters at the periphery of the viral DNA compartment. Segmentation rendering of this tomogram is illustrated in Fig. 5a and an original slice is showed in Extended Data Fig. 3f with colour overlay.

**Supplementary Movie 8 | Representative tomogram of mature virions warehouse in *B. subtilis* infected with SPP1*lacO64gp12^-^***. The tomogram exhibits the typical host cell re-organization observed upon infection with SPP1*lacO64gp12^-^*. Mature virions lacking the auxiliary capsid protein gp12, are densely packed at the periphery of the viral DNA compartment in so-called warehouses under this infection condition. Segmentation rendering of this tomogram is illustrated in Fig. 5b and an original slice is showed in Extended Data Fig. 3g with colour overlay.

**Supplementary Movie 9 | Representative tomogram of mature virions warehouse in *B. subtilis* infected with SPP1*gp6^-^***. The tomogram exhibits the typical host cell re-organization observed upon infection with SPP1*gp6^-^*. Portal-less procapsid I-like structures defective in DNA packaging and free tails accumulate in these cells. Indeed, in this condition of infection, aggregates of pre-assembled tail are found in the cytoplasm. Segmentation rendering of this tomogram is illustrated in Fig. 5c and an original slice is showed in Extended Data Fig. 3h with colour overlay.

## Notes

### Competing Interest Statement

The authors have declared no competing interest.

