## Supplementary Table 1 | Bacterial strains, viral strains, plasmids and oligonucleotides used in this work. for "Subcellular reorganization upon phage infection reveals stepwise assembly of viral particles from membrane-associated precursors"

| BIOLOGICAL RESSOURCES | SOURCE | SHORT NAME |
|---|---|---|
| **Bacterial strains** | | |
| *E. coli F- Φ80lacZΔM15 Δ(lacZYA-argF) U169 recA1 endA1 hsdR17 (rk-, mk+) phoA supE44 λ-thi-1 gyrA96 relA1* | Hanahan, 1983 | DH5α |
| *B. subtilis amyE trpC2 metB5 xin-1 attSPβ* | Yasbin et al., 1980 | YB886 |
| *B. subtilis [his met] sup$^-$ leu trpC2* | Okubo and Yanagida, 1968 | HA101B |
| *B. subtilis [metB5] sup-3 amyE trpC2 xin-1 attSPβ* | Dröge and Tavares, 2000 | BG295 |
| *B. subtilis amyE trpC2 metB5 xin-1 attSPβ (pSPW7)* | Tavares et al., 1992 | YB886 (pSPW7) |
| *B. subtilis amyE trpC2 metB5 xin-1 attSPβ thrC::($P_{pen}$-lacIΔ11-mcherry mls)* | Labarde et al., 2021 | GSY10004 |
| *B. subtilis amyE trpC2 metB5 xin-1 attSPβ thrC::($P_{pen}$-lacIΔ11-mcherry mls) sacA::(Pspac-12-mcitrine kan)* | Labarde et al., 2021 | GSY10024 |
| *B. subtilis amyE trpC2 metB5 xin-1 attSPβ thrC::($P_{pen}$-lacIΔ11-mcherry mls) sacA::($P_{spac}$-6-mcitrine kan)* | This work | GSY10027 |
| *B. subtilis amyE trpC2 metB5 xin-1 attSPβ amyE::($P_{xyl}$-rpsB-mcfp spc)* | This work | GSY10074 |
| *B. subtilis amyE trpC2 metB5 xin-1 attSPβ thrC::($P_{pen}$-lacIΔ11-mcherry mls) amyE::($P_{xyl}$-rpsB-mcfp spc)* | This work | GSY10075 |
| *B. subtilis amyE trpC2 metB5 xin-1 attSPβ thrC::($P_{pen}$-lacIΔ11-mcherry mls) sacA::($P_{spac}$-12-mcitrine kan) amyE::($P_{xyl}$-rpsB-mcfp spc)* | This work | GSY10082 |
| *B. subtilis amyE trpC2 metB5 xin-1 attSPβ amyE::($P_{xyl}$-rplA-mcfp spc)* | This work | GSY10085 |
| *B. subtilis amyE trpC2 metB5 xin-1 attSPβ thrC::(Ppen-lacIΔ11-mcherry mls) amyE ::($P_{xyl}$-rplA-mcfp spc)* | This work | GSY10086 |
| *B. subtilis amyE trpC2 metB5 xin-1 attSPβ thrC::(Ppen-lacIΔ11-mcherry mls) sacA::($P_{spac}$-12-mcitrine kan) amyE::($P_{xyl}$-rplA-mcfp spc)* | This work | GSY10087 |
| **Bacteriophage strains** | | |
| SPP1 *wild type* lytic siphophage | Riva et al., 1968 | SPP1*wt* |
| SPP1*sus70*: phage SPP1 defective in gene *1* | Behrens et al., 1979; Chai et al., 1992 | SPP1*gp1*$^-$ |
| SPP1*sus19*: phage SPP1 defective in gene *2* | Behrens et al., 1979; Chai et al., 1992 | SPP1*gp2*$^-$ |
| SPP1*sus115*: phage SPP1 defective in gene *6* | Behrens et al., 1979; Tavares et al 1992 | SPP1*gp6*$^-$ |

| | | |
|---|---|---|
| SPP1*delX110lacO64*: SPP1*delX110* derivative carrying an array of *lacO* operators | Jakutyte et al., 2011 | SPP1*lacO64* |
| SPP1*delX110lacO64sus19*: SPP1*sus19* derivative carrying deletion *delX* and an array of *lacO* operators | Labarde et al., 2021 | SPP1*lacO64gp2*⁻ |
| SPP1*delX110lacO64Δ12*: SPP1*Δ12* derivative carrying deletion *delX* and an array of *lacO* operators | Labarde et al., 2021 | SPP1*lacO64gp12*⁻ |
| SPP1*delX110lacO64sus115* : SPP1*sus115* derivative carrying deletion *delX* and an array of *lacO* operators | This work | SPP1*lacO64gp6*⁻ |
| **Plasmids** | | |
| pSPW7 | Tavares et al., 1992 | NA |
| pAL21 *bla sacA::(Pspac-mcitrine kan)* | Labarde et al., 2021 | NA |
| pAL27 *bla sacA::(Pspac-6-mcitrine kan)* | This work | NA |
| pAL29 *bla amyE::(Pxyl-mcfp spc)* | Labarde et al., 2021 | NA |
| pLG50 *bla amyE::(Pxyl-rpla-mcfp spc)* | This work | NA |
| pLG51 *bla amyE::(Pxyl-rpsb-mcfp spc)* | This work | NA |
| **Oligonucleotides** | | |
| 2677 rev: gggtgcctctctgtg | Lab collection | 2677 |
| 3185 rev: ctctggcggccacctctcgc | Lab collection | 3185 |
| 3383 fwd: gtgtccgaattgcggcccgg | Lab collection | 3383 |
| 3682 fwd: gagttgtgtggatatccaagg | Lab collection | 3682 |
| rplA_KpnI_fwd : taaccacataagga**ggtacc**tttaaaatggctaa | This work | 20 |
| rpsB_KpnI_fwd : ttagga**ggtacc**aacatgtcagtcatt | This work | 22 |
| rplA_EcoRI_rev : gg**gaattc**gccaccttttacgttaaaagttgaagagtc | This work | 28 |
| rpsB_XhoI_rev : cctttgaat**ctcgag**cgcagttgttgt | This work | 29 |
| gp6_ClaI_fwd : ggtaccgagctcgaattcaaaggagaa**atcgat**atggctgatatctacccactaggg | This work | 55 |
| gp6_XhoI_rev : gcttaccattccactgccttcgag**ctcgag**agatacttgtccagctcctcc | This work | 56 |
