## Supplementary Table 2 | Sample preparation and data collection parameters for electron cryo tomography. for "Subcellular reorganization upon phage infection reveals stepwise assembly of viral particles from membrane-associated precursors"

**Supplementary Table 2. Sample preparation and cryoET data collection parameters.**

| | **Data set 1** | **Data set 2** | **Data set 3** |
|---|---|---|---|
| **Samples with number of tilt-series analysed** | Uninfected (9), infection with SPP1*gp12*⁻ (9) | Uninfected (18); infection with SPP1*wt* (75), SPP1*gp1*⁻ (9), SPP1*gp2*⁻ (28), SPP1*gp6*⁻ (58) | Infection with SPP1*wt* (9); SPP1*gp12*⁻ (16) |
| Grid type | Quantifoil R2/2 Cu 200 | | |
| Grid glow discharge | 30 sec - 15 mA (Pelco easiGlow™) | | |
| Vitrification | Leica GP | | |
| Sample Volume | 4 µL | | |
| Sample concentration | $10^9$ cells/mL | | |
| Blotting position | 44 and 4.4 | | |
| Blot paper | Whatman 1 | | |
| Blot time | 20 sec | | |
| Temperature | 22 °C | | |
| Relative humidity | 80 % | | |
| Wait time | 0 sec | | |
| Cryogen | Ethane | | |
| Microscope | Titan Krios G3 | | |
| Voltage | 300 kV | | |
| Energy filter / Detector | Gatan BioQuantum-K3 | | |
| Stilt width | 20 eV | | |
| Objective aperture | 70 µm | | |
| Nominal Magnification | 26,000X | 42,000X | 64,000X |
| Pixel size | 3.336 Å | 2.15 Å | 1.372 Å |
| Defocus range | -4 to -5 µm | | |
| Acquisition Scheme | Dose-symmetric tilt scheme: +/- 60 degrees ; 3 degree increment | | |
| Dose per frame | 5e-/Å$^2$ over 3 frames | 4.5 e-/Å$^2$ over 5 frames | 2.5e-/Å$^2$ over 2 frames |
| Total dose | ~200 e-/Å$^2$ | ~180 e-/Å$^2$ | ~120 e-/Å$^2$ |
